# Multivalent interactions of the disordered regions of XLF and XRCC4 foster robust cellular NHEJ and drive the formation of ligation-boosting condensates *in vitro*

**DOI:** 10.1101/2023.07.12.548668

**Authors:** Duc-Duy Vu, Alessio Bonucci, Manon Brenière, Metztli Cisneros-Aguirre, Philippe Pelupessy, Ziqing Wang, Ludovic Carlier, Guillaume Bouvignies, Patricia Cortes, Aneel K. Aggarwal, Martin Blackledge, Zoher Gueroui, Valérie Belle, Jeremy M. Stark, Mauro Modesti, Fabien Ferrage

## Abstract

In mammalian cells, DNA double-strand breaks are predominantly repaired by non-homologous end joining (NHEJ). During repair, the Ku70/80 heterodimer (Ku), XRCC4 in complex with DNA Ligase 4 (X4L4), and XLF form a flexible scaffold that holds the broken DNA ends together. Insights into the architectural organization of the NHEJ scaffold and its regulation by the DNA-dependent protein kinase catalytic subunit (DNA-PKcs) have recently been obtained by single-particle cryo-electron microscopy analysis. However, several regions, especially the C-terminal regions (CTRs) of the XRCC4 and XLF scaffolding proteins, have largely remained unresolved in experimental structures, which hampers the understanding of their functions. Here, we used magnetic resonance techniques and biochemical assays to comprehensively characterize the interactions and dynamics of the XRCC4 and XLF CTRs at atomic resolution. We show that the CTRs of XRCC4 and XLF are intrinsically disordered and form a network of multivalent heterotypic and homotypic interactions that promotes robust cellular NHEJ activity. Importantly, we demonstrate that the multivalent interactions of these CTRs led to the formation of XLF and X4L4 condensates *in vitro* which can recruit relevant effectors and critically stimulate DNA end ligation. Our work highlights the role of disordered regions in the mechanism and dynamics of NHEJ and lays the groundwork for the investigation of NHEJ protein disorder and its associated condensates inside cells with implications in cancer biology, immunology and the development of genome editing strategies.

## INTRODUCTION

DNA double-strand breaks (DSBs) can be induced by external agents such as ionizing radiation or by internal processes, including errors in DNA transactions and the action of reactive oxygen species produced by oxidative metabolism (1, 2). DSBs also form as intermediates during programmed genome rearrangements in developing lymphocytes (3). If undetected or mis-repaired, DSBs threaten genomic integrity as they may lead to chromosomal rearrangements that can trigger tumor formation or cell death. Several pathways have evolved in eukaryotic cells to mend DSBs, among which the non-homologous end joining (NHEJ) pathway is prevalent (1, 4, 5).

NHEJ is a flexible process both in terms of structures and mechanisms (1, 4, 5). NHEJ is rapid and adaptable to repair any type of broken DNA end structures. This is achieved by the assembly of an untemplated synaptic scaffold that brings and holds together the two free-broken DNA ends, recruits end-processing factors, and ultimately promotes the ligation step (1, 4, 5). The dynamic nature and the involvement of numerous effectors of NHEJ render the investigation of this mechanism challenging, especially in determining how the core effectors assemble and coordinate to ensure effective DNA repair.

Many contributions from structural and single-molecule studies have yielded clear pictures of several essential steps in the NHEJ pathway. In the consensus model, the Ku70/80 heterodimer (Ku) initially threads onto broken DNA ends and directly attracts the DNA-dependent protein kinase catalytic subunit (DNA-PKcs), XLF, and XRCC4 in complex with DNA Ligase 4 (X4L4) to assemble into a synaptic complex that tethers and re-ligates DNA ends (1, 4–13). Single-particle cryo-electron microscopy (cryo-EM) studies of NHEJ supercomplexes containing Ku, XLF, XRCC4, and DNA Ligase 4 (LIG4) with or without DNA-PKcs assembled on DNA have revealed several structural organizations of the NHEJ synaptic scaffold and suggested how transitions between different states may occur (11, 14–17). However, putatively disordered C-terminal regions (CTRs) of both XLF and XRCC4 are mostly unresolved, likely due to their heterogeneous conformations under cryogenic condition.

The CTRs of XLF and XRCC4 have been considered dispensable for NHEJ, but recent studies suggest otherwise (17–19), even though their functional role during NHEJ remains undefined. Here, we use cellular models to show that the CTRs of XRCC4 and XLF are crucial for robust NHEJ activity. We use nuclear magnetic resonance (NMR) to understand their functional role at atomic resolution. We demonstrate that both CTRs of XLF and XRCC4 are intrinsically disordered and form a network of weak multivalent interactions, which drives the formation of condensates of XLF and X4L4 *in vitro*. XLF and X4L4 condensates recruit other NHEJ factors and strongly stimulate DNA end ligation, offering a yet unexplored perspective on the organization in space and time of the DSB repair mechanism by NHEJ.

## RESULTS

### The CTRs of XLF and XRCC4 promote robust cellular NHEJ

XLF and XRCC4 share a similar structural organization, including a head domain (Head), a coiled-coil domain (CC), and a CTR (**Fig. 1a-b)** (20, 21). To test the importance of the CTRs of XLF and XRCC4 for cellular NHEJ, the EJ7-GFP reporter was used to measure end joining between distal ends of two Cas9-mediated blunt chromosomal DSBs without indel mutations from the edge of the DSB (**Fig. 1c**). Such No Indel end joining causes restoration of a GFP+ cassette that is measured by flow cytometry, and this repair event has previously been shown to require both XRCC4 and XLF (22). To examine mutations in both factors simultaneously, we used a previously described XRCC4-KO/XLF-KO HEK293 cell line with a EJ7-GFP reporter integrated in the genome (23), which was transfected with the plasmids for Cas9/sgRNAs to induce DSBs, along with expression vectors for XLF and XRCC4, and the truncated mutants XLF_1-251_ and XRCC4_1-230_. For XLF, we also combined the truncation mutation with L115A, which weakens the binding interface with XRCC4 (20, 22, 24, 25). Of note, all XRCC4 and XLF expression constructs express untagged versions of the proteins. We found that combining the deletion of both CTRs with the XLF_1-251_ L115A mutant and XRCC4_1-230_ caused a marked (9.8-fold) decrease in No Indel end joining, compared to XRCC4_WT_ (**Fig. 1d-e**).

**Figure 1.**
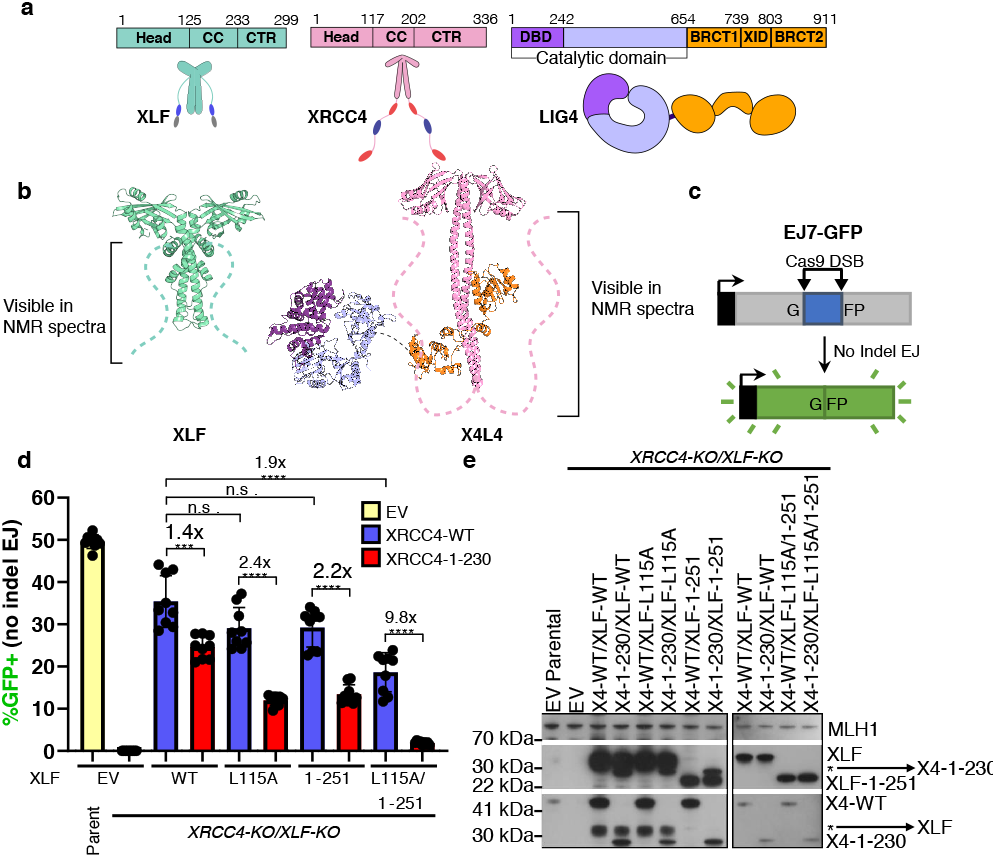
The CTRs of XLF and XRCC4 promote robust cellular NHEJ. (**a**) Cartoons of the XLF, XRCC4 and LIG4 proteins with annotated domains. XLF and XRCC4 form homodimers. Head: head domain; CC: coiled-coil, CTR: C-terminal region, DBD: DNA-binding domain, XID: XRCC4-interaction domain; BRCT1/BRCT2: first/second BRCA1 C-terminal domain (**b**) The crystal structure of XLF (PDBid 2R9A) and the Cryo-EM structure of XRCC4 in complex with LIG4 (X4L4) (PDBid 7LSY).The dashed lines represent the CTRs which are lacking in these structures but are visible in NMR spectra in **Fig. 2a** and **Fig. 3a** for XLF and XRCC4, respectively. (**c**) The EJ7-GFP reporter (not to scale) for No Indel EJ between two Cas9/sgRNA DSBs, which was integrated into XRCC4-KO/XLF-KO HEK293 cells. (**d**) GFP frequencies are normalized to parallel GFP transfections to account for transfection efficiency. (**e**) Immunoblots show levels XRCC4_WT_, XRCC4_1-230_, XLF_WT_, and XLF mutants, *indicates background XLF or XRCC4. n = 9 biologically independent transfections. Statistics with unpaired two-tailed t-test with Holm-Sidak correction. #x represents fold effect. ****P<0.0001, ***P<0.001, **P<0.01, n.s. = not significant. Data are represented as mean values *±* SD.

The XRCC4_1-230_ mutant also showed a significant but modest decrease vs. XRCC4_WT_ when combined with either XLF, XLF_L115A_, or XLF_1-251_ (1.4, 2.4, and 2.2-fold, respectively, **Fig. 1d**). Similarly, the XLF_1-251_ L115A mutant showed a modest decrease vs. XLF_WT_ when combined with XRCC4_WT_ (1.9-fold). Expression levels of the XRCC4_1-230_ mutant appears to be less than for XRCC4_WT_ (**Fig. 1e**), however both are higher than endogenous XRCC4, with which the XLF_1-251_ L115A mutant only causes a 1.3-fold decrease (**Fig. S1**), which is much smaller than the 9.8-fold decrease when combined with XRCC4_1-230_ (**Fig. 1d**). We conclude that the CTRs of XLF and XRCC4 synergize to promote robust cellular NHEJ. We present below our NMR investigation of the CTRs of XLF and XRCC4 at the atomic resolution to decipher their functional roles.

### Interactions of the disordered CTR of XLF

The CTR of XLF is predicted to be disordered (**Fig. S2c**). To confirm the disordered nature of this region with NMR spectroscopy, isotopically labeled full-length XLF and its CTR in isolation (residues 229-299, XLF_CTR_) were produced. Despite the large molecular weight of the XLF homodimer (*¥* 66 kDa), the ^1^H-^15^N HSQC correlation spectrum of isotopically labeled full-length XLF showed sharp and well-resolved peaks (**Fig. 2a**). Most peaks of the spectra of full-length XLF and of the XLF_CTR_ alone are superimposed. We used a series of triple-resonance NMR experiments to assign backbone resonances of all non-proline residues the CTR, between residues 232 and 299 (**Fig. S3a**). ^1^H chemical shift dispersion, secondary structural propensity, ^15^N-{^1^H} nuclear Overhauser effects (hNOE), and small-angle X-ray scattering experiment jointly indicated the lack of stable secondary structure elements in the XLF_CTR_ either alone or in the context of the full-length protein construct (**Fig. S4a**,**b, S5**), defining it as an intrinsically disordered region (IDR).

**Figure 2.**
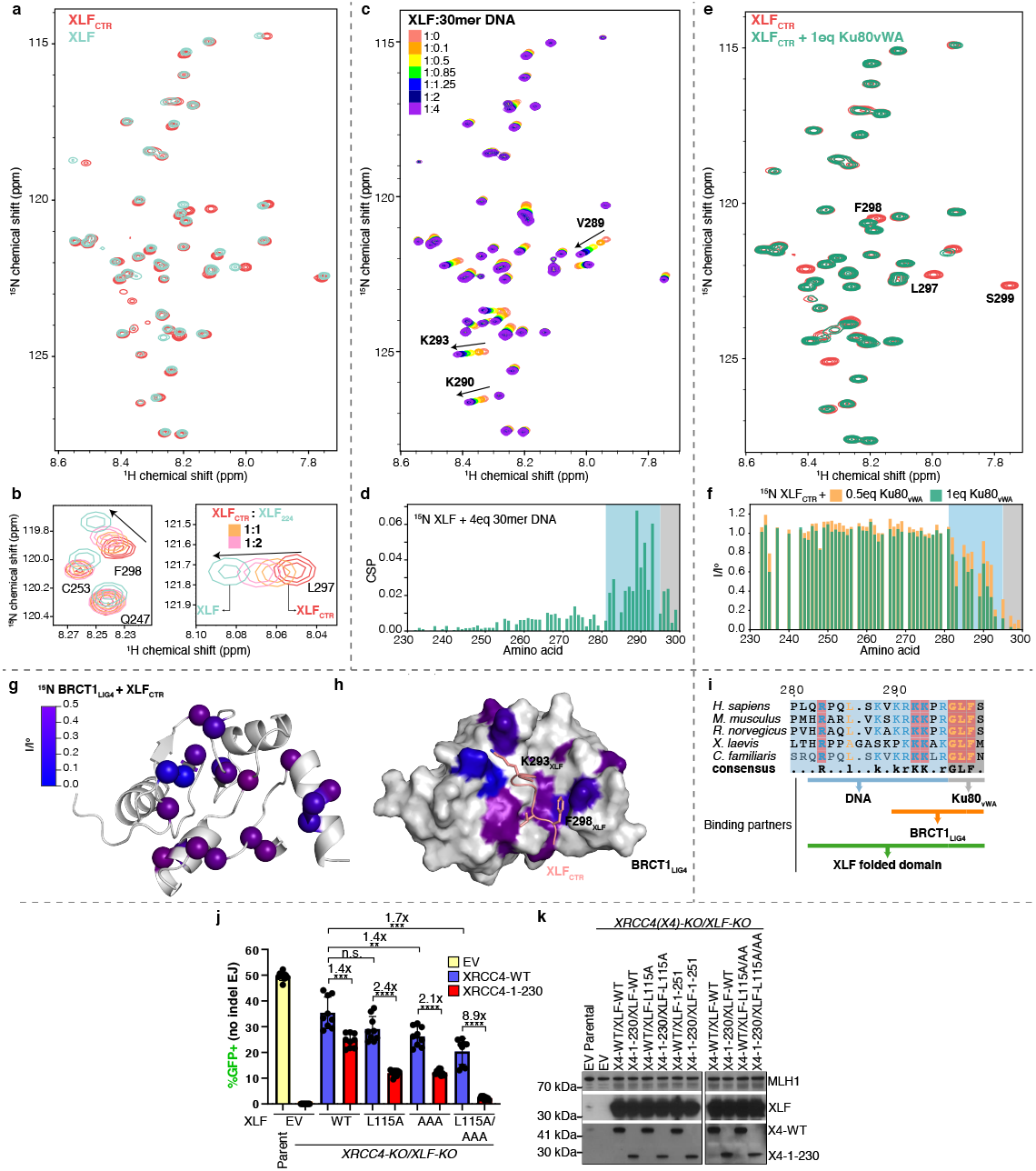
Multivalent interactions within the CTR of XLF. (**a**) Overlay ^1^H-^15^N HSQC spectra of full-length XLF (cyan) and XLF_CTR_ (red). (**b**) Insets from superimposed ^1^H-^15^N HSQC spectra of 250 *μ*M XLF_CTR_ (red) after adding one (orange) or two (pink) equivalents of XLF_1-224_ compared with the spectrum of full-length XLF (cyan). (**c**) Overlay ^1^H-^15^N HSQC spectra of 150 *μ*M XLF recorded in a titration with 30mer DNA. (**d**) Chemical shift perturbations (CSP) of XLF upon addition of four equivalents of 30mer DNA. (**e**) Overlay of ^1^H-^15^N HSQC spectra of 50 *μ*M XLF_CTR_ alone (red) and in the presence of one equivalent of Ku80_vWA_ (green). (**f**) Peak intensity ratios from ^1^H-^15^N HSQC spectra of ^15^N XLF_CTR_ spectra comparing the intensity in the spectrum of XLF_CTR_ alone (I°) and after adding 0.5 (orange) or 1 (green) equivalent of Ku80_vWA_. (**g**) The residues of BRCT1 that experienced significant peak intensity reduction (I/I° < 0.5) after adding XLF_CTR_ are mapped as a color gradient (0: blue, 0.5: purple) on the structure of BRCT1 (generated by Alphafold-multimer). I/I°: normalized peak intensity of BRCT1 before and after adding two equivalents of XLF_CTR_. (**h**) The model of BRCT1-XLF_CTR_ complex predicted by Alphafold-multimer (26), the affected residues are color-mapped as in (**g**). (**i**) Sequence alignment of the last 20 residues of XLF homologs. Conserved residues are colored with the following scheme: hydrophobic residues and glycine, orange; positively charged residue, blue; strictly conserved residues are highlighted in red. The residues of the CTR of XLF that interact with different partners are indicated. (**j**) Combining loss of the CTR of XRCC4, along with the XLF_AAA/L115A_ mutant causes a marked decrease in cellular NHEJ. GFP frequencies are normalized to parallel GFP transfections to account for transfection efficiency. n = 9 biologically independent transfections. The values of XLF_EV_, XLF_WT_, and XLF_L115A_, with or without XRCC4_WT_ or XRCC4_1-230_ are the same as those in **1d**. Statistics with unpaired two-tailed t-test with Holm-Sidak correction. #x represents fold effect. ****P<0.0001, ***P<0.001, **P<0.01, n.s. = not significant. (**k**) Immunoblots show levels XRCC4_WT_, XRCC4_1-230_, XLF_WT_, and XLF mutants. Data are represented as mean values *±* SD.

The NMR spectra of the disordered CTR within the full XLF homodimer, make it possible to characterize interactions of the CTR, which were previously explored either indirectly or with reductions to short peptides (27–29). The interaction of XLF and Ku is well known (27–29). The Ku binding motif (KBM) of XLF has been defined as the last ten residues of XLF (27) and the structures of Ku bound to XLF_KBM_ peptide have been solved by X-ray crystallography (28). We titrated ^15^N XLF_CTR_ with the Ku80 von Willebrand domain (Ku80_vWA_) and observed that XLF-Ku interactions mainly involved the last five residues (^295^RGLFS^299^), while residues 283-294 were only slightly perturbed upon binding to Ku80_vWA_ (**Fig. 2e-f**), in agreement with the crystal structure of Ku with XLF_287-299_ peptide where only the residues ^296^GLFS^299^ occupy the hydrophobic pocket of Ku80_vWA_ to stabilize the XLF-Ku interaction (28). This led us to define the core KBM of XLF as residues R295-S299.

Next, we characterized the interaction of XLF with DNA. The CTR of XLF was shown to play an important role in the DNA-binding capacity of the full-length protein (30, 31). Yet, the region of interaction was not clear (32). We monitored chemical shift perturbations (CSPs) of XLF upon the addition of DNA to better characterize the region of interaction of XLF with DNA (**Fig. 2c,d**, **S6**). The presence of stoichiometric amounts of double-stranded DNA induced significant CSPs in the ^1^H-^15^N HSQC correlation spectrum of XLF. The largest perturbations were observed for residues R283, V289, K290, R291, and K293, whereas smaller perturbations were detected for the residues in the KBM, suggesting that this motif does not significantly contribute to DNA binding. Consistently, the K293A mutation greatly reduced the affinity of XLF for DNA, while the K290/R291A double mutation abolished the interaction. In contrast, the L297A and F298A mutations in the KBM did not significantly affect the interaction (**Fig. S6d**). Therefore, the positively charged enriched region (residues 282-294) is the core of the DNA-binding motif (DBM) (**Fig. 2i**).

XLF binds DNA in a length-dependent manner (31, 33). The region 290-296 of XLF shows sequence similarities with AT-hook motifs that preferentially bind to the minor groove of AT-rich sequences of double-stranded DNA (34). To decipher the binding mode of XLF to DNA, XLF was titrated with double-stranded DNA of different lengths (15mer, 18mer, 21mer, 30mer) and sequence compositions (AT-rich, GC-rich and random sequence) (**Fig. S6**). The apparent dissociation constants (K_D_) were quantified by fitting the 2D lineshapes and positions of the affected peaks with a two-state binding model (35). The apparent K_D_ values of XLF for double-stranded DNA with 21mer and 30mer ranged from 4 to 16 μM and increased significantly to 34 and 141 μM for 18mer and 15mer DNA, respectively. This is in agreement with the length-dependent affinity of XLF for DNA, as observed in previous studies (31, 33). XLF had comparable affinities for AT-rich, GC-rich or random balanced sequences (**Fig. S6**), despite the similarity of the DBM of XLF to AT-hook motifs (34). To identify the binding mode of the CTR of XLF to DNA, we carried out a competitive binding experiment between XLF, AT-rich DNA and Netropsin (**Fig. S7**), a small molecule that binds specifically to the minor groove of AT-rich DNA (36). Netropsin competed efficiently with XLF on AT-rich DNA, suggesting that XLF and Netropsin share the same binding site. Thus, the residues 282 to 295 in the CTR of XLF bind to DNA, most likely in the minor groove, in a length-dependent and sequence non-specific manner.

Next, we identified the presence of homotypic interactions in full-length XLF that involve the CTR. The backbone ^15^N transverse relaxation rates (R_2_) in the DBM and KBM of the full length XLF exhibited significantly elevated values compared to the XLF_CTR_ alone (**Fig. S8a**), suggesting transient interactions with the folded domain. Accordingly, adding the folded domain of XLF (XLF_1-224_) to a solution of ^15^N XLF_CTR_ increased the values of R_2_ for the residues in the DBM and KBM similarly as in full-length XLF. In addition, ^1^H and ^15^N chemical shifts of residues L297 and F298 shifted back towards the values in full-length XLF (**Fig. 2b**). These results demonstrate that the DBM and KBM of XLF_CTR_ interact with the folded domain of XLF. Moreover, the KBM and DBM of XLF_CTR_ interacted synergistically with the folded domain of XLF, as the interaction was abolished only when key residues in both DBM (K290/R291A) and KBM (F298A) were mutated to alanine (**Fig. S8b**). We also observed that a non-cleavable hexahistidine tag at the C-terminus of XLF, as used in several previous studies (19, 30), severely alters the R_2_ profile of XLF_CTR_ (**Fig. S3b**), possibly by artificially stabilizing these intramolecular interactions.

Where do the DBM and KBM bind onto the folded domain of XLF? The large molecular weight of the XLF dimer and the possible formation of non-specific oligomers at high concentrations (20) prevented us from directly observing the folded domain in NMR spectra. As an alternative strategy, paramagnetic relaxation enhancement (PRE) experiments were used to identify the binding sites of DBM and KBM onto the folded domain of XLF (**Fig. S9**). In these experiments, a paramagnetic probe (a nitroxide) was attached to a cysteine side chain from the folded domain of a single-cysteine mutant of XLF. The strong interaction between protons and the electron spin leads to decreased intensity in the ^1^H-^15^N NMR spectrum for residues that are in the proximity, even transiently, of the nitroxide probe, typically within 15 Å (37, 38). Peaks from the KBM motif lost intensity when the probe was attached to the head domain (C74), while peaks from the DBM weaken when the probe was attached to the coiled-coil domain (C188) (**Fig. S9**). These PRE patterns corroborate our hypothesis that the DBM and the KBM of XLF_CTR_ interact with the folded domain at distinct binding sites.

We next investigated the role of the CTR of XLF in the interaction with the X4L4 complex. In pull-down assays, XLF interacted with LIG4 through the first BRCT and XRCC4-interacting domains (BRCT1-XID) (39), but the binding site remains unknown. XLF forms oligomers with XRCC4 and X4-BRCTs through XLF-XRCC4 head-to-head interaction (30) as confirmed by ^15^N-edited diffusion measurements in solution (**Fig. S10f**). Importantly, severe intensity losses were observed for the KBM and DBM of XLF upon the addition of X4-BRCTs (**Fig. S10a**). Signals for the KBM and DBM were attenuated even when the XLF-XRCC4 head-to-head interaction was disrupted by the L115D mutation (25) (**Fig. S10b**,**c**) but not when the BRCT1 domain was removed (**Fig. S10d**,**e**). Taken together, these data indicate a direct interaction between the KBM and DBM of XLF and the BRCT1 domain of LIG4.

To identify the XLF-binding region in the LIG4 BRCT1, the non-proline backbone chemical shifts of the LIG4 BRCT1 domain (LIG4 residues 654-759) were assigned (**Fig. S11a**). Adding the XLF_CTR_ to isotopically labeled BRCT1 led to a reduction of peak intensities and changes in chemical shifts (**Fig. 2g, S11b**,**c**). Using AlphaFold-multimer (26), a model of the BRCT1-XLF_CTR_ complex was computed (**Fig. S11d**). Although the predicting confidence for XLF_CTR_ is low, the predicted interface agrees with the perturbation in the NMR spectra for both proteins. In this model for the molecular basis of the XLF_CTR_-BRCT1 interactions (**Fig. 2h**), residue F298 of XLF forms hydrophobic interactions with residues H753 and F754, whereas K293 of XLF further stabilizes the complex through electrostatic interactions with residue D661 of LIG4. The results also explain why the CTR of XLF can indirectly stabilize the interaction between XLF and XRCC4 as observed previously (18, 40). Altogether, we show that the disordered CTR of XLF is an interaction hub, which binds weakly to Ku, DNA, LIG4 and the folded domain of XLF in an intra-or intermolecular fashion.

To probe the role of specific interaction of the CTR of XLF for cellular NHEJ activity, we designed a triple mutant of XLF, with the mutations K290A and R291A in the DBM as well as F298A in the KBM (XLF_AAA_) that impaired the interaction of the CTR of XLF with DNA (**Fig. S6d**), the folded domain of XLF (**Fig. S8b**), Ku80_vWA_ domain (**Fig. 2f**) (28) and potentially the LIG4 BRCT1 domain (**Fig. 2g-h**). When tested for its ability to support NHEJ in cells with the EJ7-GFP reporter, the XLF_AAA_ mutant behaved similarly to the XLF_1-251_ truncation described above, and notably caused a marked (8.9-fold) decrease in No Indel end joining when combined with L115A and with XRCC4_1-230_ (**Fig. 2j-k**). These results highlight the key role of these specific interactions of the CTR of XLF for robust NHEJ activity in cells.

### Interactions of the disordered CTR of XRCC4

The CTR of XRCC4 is longer than the CTR of XLF, encompassing residues 202-336 and is predicted to be disordered (**Fig. S2c**). Low-resolution cryo-EM structure classification (41) suggests that the CTR of XRCC4 may fold back to the coiled-coil domain. To characterize the structural properties of the CTR of XRCC4, we used isotopically labeled full-length XRCC4 and a protein fragment corresponding to its CTR (XRCC4_CTR_). Two additional protein samples containing isotopically labeled XRCC4 in complex with full-length LIG4 (X4L4) or the unlabeled BRCT tandem of LIG4 (X4-BRCTs) were also studied. The ^1^H-^15^N HSQC spectrum of XRCC4 and XRCC4_CTR_ displayed low proton chemical shift dispersion, a signature of disordered protein regions (**Fig. 3a, S3c-e**), with no detectable signal from the folded domain (42). Inspection of the ^1^H-^15^N HSQC spectra of all the constructs and complexes identified only small differences, indicating no major conformational changes in the CTR of XRCC4. All non-proline residues from 208-366 were assigned unambiguously except residues 273 to 278. (**Fig. S3c**). NMR relaxation and chemical-shift analyses together with SAXS experiments confirmed the intrinsically disordered nature of the CTR of XRCC4 across all the constructs (**Fig. S4c**,**d, S12**). In addition, continuous-wave electron paramagnetic resonance (CW-EPR) spectra also showed that the binding to LIG4 BRCTs did not significantly alter the dynamic properties of the CTR of XRCC4 (**Fig. 3b**).

**Figure 3.**
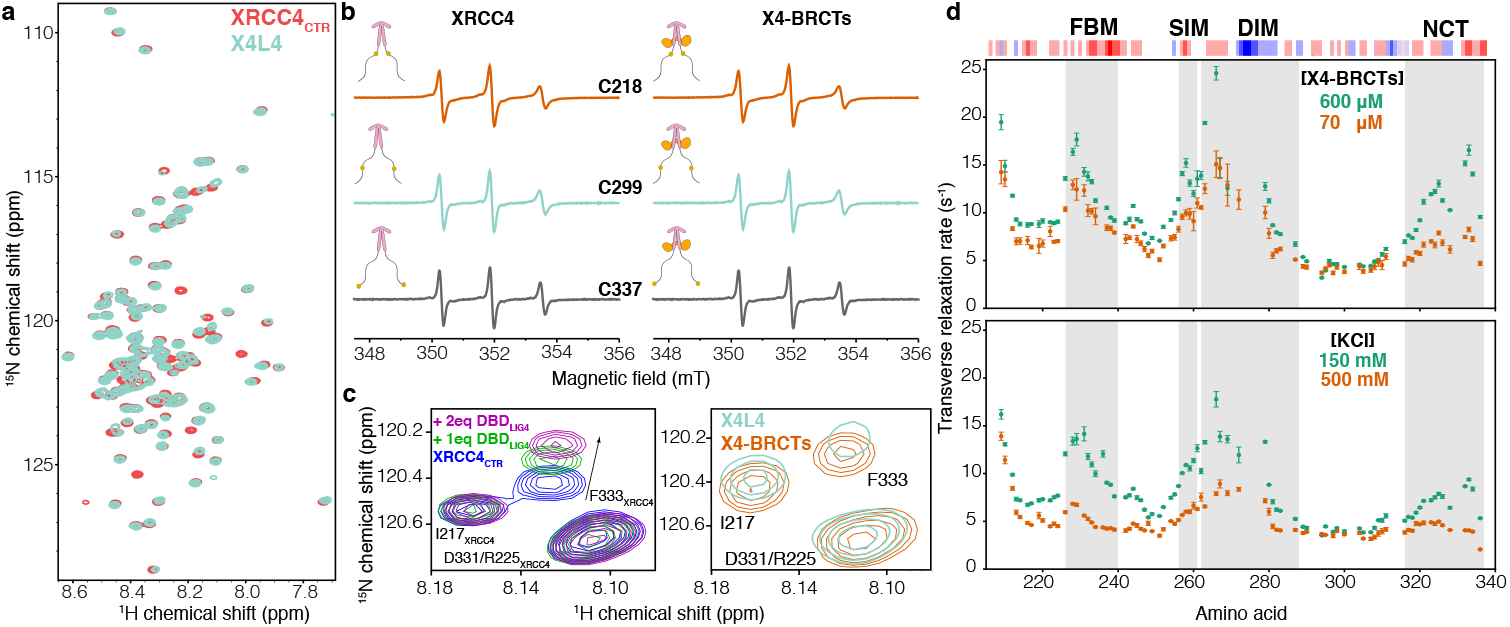
Multivalent interactions within the CTR of XRCC4. (**a**) Overlay of ^1^H-^15^N HSQC spectra of XRCC4_CTR_ (red) and XRCC4 in complex with LIG4 (X4L4, cyan). (**b**) CW-EPR spectra (X-band) recorded at room temperature for XRCC4 variants C218-MTSL (orange), C299-MTSL (cyan), and C337-MTSL (gray). The simulated spectra and the simulation parameters adopted for each spectrum are reported in **S21**). (**c**) Left: the inset shows a part of the superimposed spectra of ^15^N XRCC4_CTR_ upon adding 1 (green) or 2 (pink) equivalents of DBD_LIG4_; right: the inset shows the same region of the superimposed spectra of ^15^N full-length XRCC4 in complex with LIG4 (cyan) or BRCT tandem (X4-BRCTs, orange), the shift of the peak corresponding to the residue F333 of XRCC4_CTR_ is highlighted with the arrow. (**d**) *upper panel*: The transverse relaxation rate (R_2_) of backbone ^15^N nuclei for residues corresponding to the XRCC4_CTR_ in the X4-BRCTs complex at concentrations 600 and 70 μM; *lower panel*: the R_2_ of backbone ^15^N nuclei for XRCC4 in the X4-BRCTs complex (300 μM) with KCl concentrations 150 mM (green) and 500 mM (orange). The R_2_ were measured at 298 K on the 800 MHz spectrometer. The four regions (FBM, SIM, DIM, and NCT) are highlighted in gray; the three-residue rolling charge distribution is displayed on top of the graph.

Based on sequence conservation and functional roles (**Fig. S2d**), four subregions of XRCC4_CTR_ were identified: the Forkhead-associated-domain Binding Motif (FBM, residues 226-240) (43); the SUMO Interaction Motif (SIM, residues 256-261) (23); the DNA-PKcs Interaction Motif (DIM residues 262-288) (11, 17); the Negatively Charged C-terminal region (NCT, residues 316-336) (**Fig. 3d, S2d**). When titrating the XRCC4_CTR_ with 21mer double-stranded DNA, small changes in the chemical shift of residues within the DIM region were observed, suggesting that this region can bind transiently to DNA (**Fig. S13a, b)**. This is in line with our previous results showing the CTR of XRCC4 does not contribute significantly to the DNA-binding capacity of the full-length protein (44) but is important for the DNA-bridging property of XLF-XRCC4 complexes (19).

The four conserved regions within the CTR of XRCC4 exhibited high backbone ^15^N transverse relaxation rates R_2_, indicating the likely presence of local interactions (**Fig S12a**,**f**). The R_2_ relaxation rates increased with increasing concentration (**Fig. 3d**_*upper panel*_), suggesting the presence of intermolecular interactions between XRCC4 dimers mediated by the CTR of XRCC4. Considering the uneven charge distribution in XRCC4_CTR_, we hypothesized that those interactions were driven by electrostatics. Indeed, increasing salt concentrations significantly decreased the R_2_ value, indicating weakened interactions (**Fig. 3d**_*lower panel*_). To identify specific long-range interactions between the CTR of XRCC4 subregions, PRE experiments on X4-BRCTs complexes were carried out (**Fig. S14**). Comparison of the experimental PREs with the results calculated from random unconstraint conformations revealed that the CTR of XRCC4 is not fully extended but that its four subregions interact with one another. To test whether these “long-range” interactions within the CTR of XRCC4 are inter-or intra-monomer, we measured the PRE effects for the XRCC4_CTR_ alone (**Fig. S15**). Long-range interactions involving the four conserved regions were still observed with the XRCC4_CTR_ fragment but to a lesser extent. To directly characterize the conformational space of XRCC4_CTR_, the ensemble of XRCC4 was generated using flexible-meccano and the ASTEROIDS algorithm (45–47) with backbone chemical shifts, intramolecular PREs, and SAXS data (**Fig. S16**). The XRCC4_CTR_ ensemble clearly showed a lack of significantly populated secondary structure, with only slightly elevated alpha-helical propensity in the NCT subregion, and the presence of long-range contacts between the SIM-DIM subregions and the FBM-NCT subregions. In brief, these data demonstrate that the four subregions within the CTR of XRCC4 can form a network of electrostatic interactions within the same monomer, between two monomers within an XRCC4 dimer, and between XRCC4 dimers.

Finally, we identified that the CTR of XRCC4 interacts with the DNA binding domain (DBD) of LIG4. The ^1^H-^15^N HSQC signals from XRCC4 residues around F333 exhibited significant chemical shift and intensity perturbations between X4-BRCTs and X4L4 (**Fig. 3c**), suggesting an interaction of the CTR of XRCC4 with the catalytic or DNA binding domains of LIG4. Accordingly, titration of ^15^N XRCC4_CTR_ with unlabeled DBD of LIG4 reproduced the perturbations around the residue F333 as observed in X4L4 (**Fig. 3c, S13c**,**d**), demonstrating the interaction between the CTR of XRCC4 and the DBD of LIG4.

### Liquid-liquid phase separation of XLF and XRCC4 in complex with LIG4

Networks of weak multivalent interactions involving IDRs have been proposed to facilitate protein-protein and protein-nucleic acid interactions to increase local concentrations leading to the formation of condensates, which can, in turn, undergo liquid-liquid phase separation (LLPS) (48–50). The formation of protein or protein/RNA condensates has been observed at DNA repair sites inside cells (51–53), but LLPS of NHEJ proteins has not been reported yet. We explored whether the many multivalent interactions involving the CTRs of XLF and XRCC4 could induce LLPS. In the presence of Ficoll 400 as a crowding agent, X4L4 co-phase-separated with XLF at concentrations as low as 500 nM and did not require DNA to undergo LLPS (**Fig. 4a**,**b, S17**). However, DNA was able to partition into XLF and X4L4 droplets (**Fig. 4a**). The dense protein phase showed liquid properties as evidenced by the observation of droplet-like fusion events and partial exchange (around 20%) with the surrounding environment in fluorescence recovery after photobleaching (FRAP) experiments (**Fig. 4c**,**d**).

**Figure 4.**
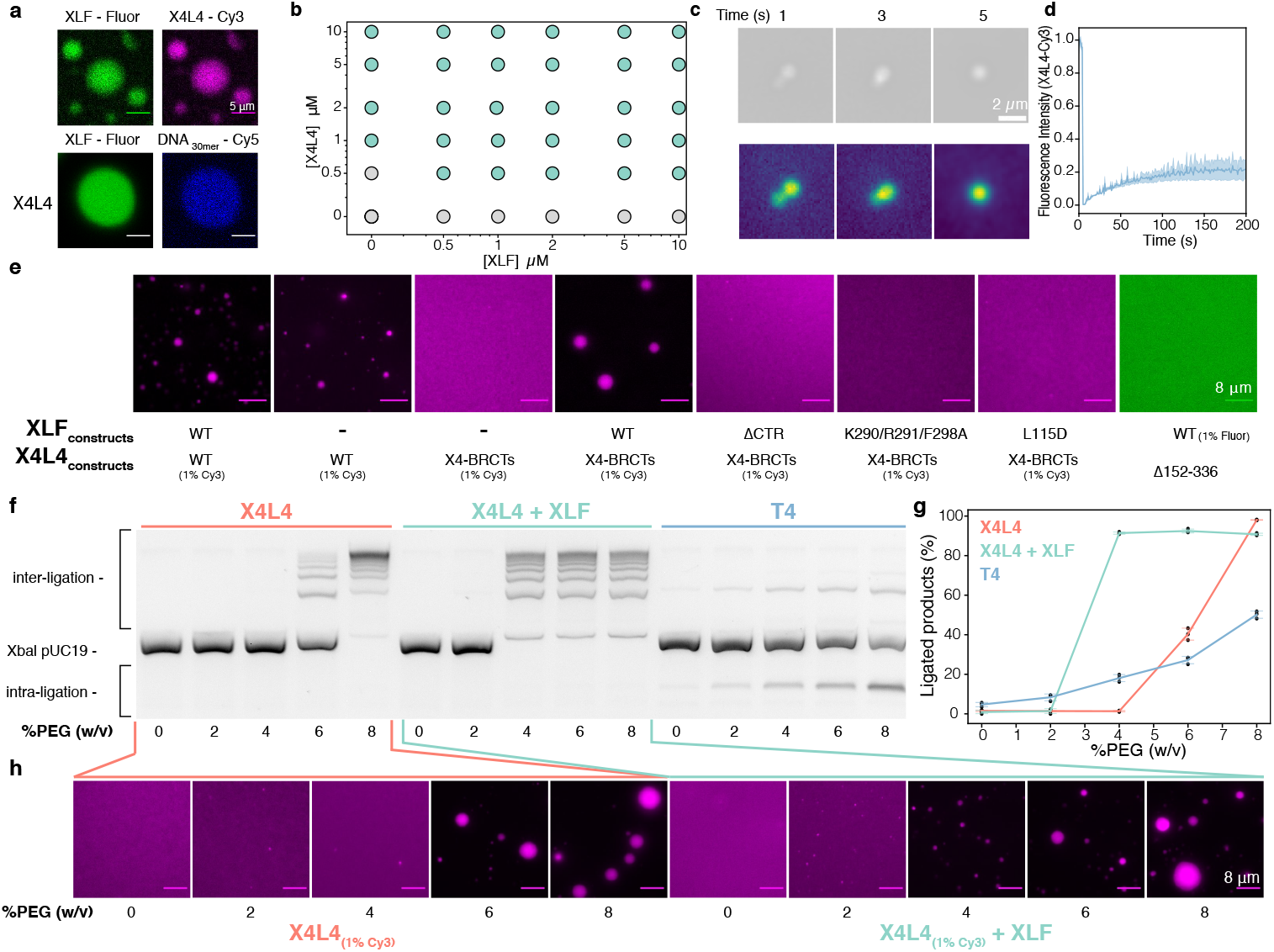
X4L4 and XLF undergo liquid-liquid phase separation *in vitro*. (**a**) Co-localization of XLF with X4L4 and DNA within the droplets observed by the fluorescence microscopy with Fluorescein-labeled XLF, Cy3-labeled X4L4, and Cy5-labeled DNA, the scale bar is 5 *μ*m. (**b**) Phase diagram of XLF with X4L4, the grey and green circles represent monophasic and biphasic solutions, respectively. (**c**) Droplet fusion event of X4L4 and XLF droplets (top), the same images are plotted as intensity gradient to highlight the contrast. (**d**) FRAP experiments of X4L4-XLF droplets, the experiments were carried out as duplicate. (**e**) Ability of different XLF and X4L4 constructs to undergo LLPS. The concentrations of all constructs were fixed at 10 μM, refer to (**Fig. S19, S20**) for a complete range of concentrations. (**f**) Image of an agarose gel after fractionation by electrophoresis and SYBR Green I staining of deproteinized cohesive end ligation products from reactions containing X4L4, X4L4-XLF, or T4 with an increasing amount of PEG8000. The substrate was 2.7 kb XbaI-linearized pUC19 plasmid. The reaction time was 5 min. (**g**) Quantification of ligated DNA of the reactions is shown on the left, the experiments were carried out in triplicate. (**h**) The same reactions as in (**f**) with 1% Cy3 labeled X4L4 were visualized using fluorescence microscopy. Data are represented as mean values ± SD.

Next, we sought to determine the minimal components required for LLPS of XLF and X4L4. X4L4 alone can undergo phase transition but requires higher concentrations of X4L4 (around 1 μM) (**Fig. 4b**). Removing the catalytic domain of LIG4 from the X4L4 complex (X4-BRCTs) impaired its ability to form droplets possibly due to the interaction between the CTR of XRCC4 and the DBD of LIG4 (**Fig. S18**). The XRCC4 phospho-mimetic mutant in complex with the LIG4 BRCTs (19) did not undergo LLPS (**Fig. S18**), but adding XLF to this X4-BRCTs complex restores phase separation at an increased critical concentration of 2 μM (**Fig. S18, S19**).

To determine the molecular interactions that govern LLPS, a series of XLF and XRCC4 mutants were investigated (**Fig. 4e, S19, S20**). Truncating the CTR of XLF or introducing the triple mutation (K290/R291/F298A) impaired the formation of XLF and X4-BRCTs droplets, most likely due to the disruption of the interactions of the CTR of XLF with the folded domain of XLF and with the BRCT1 of LIG4 (**Fig. 2i**). In contrast, mutations in either the DBM (K293A or K290R291A) or the KBM (L297E or F298A) alone did not fully prevent phase separation. Disrupting the interaction between XLF and XRCC4 by introducing the L115D mutation in XLF abolished LLPS.

For XRCC4 and LIG4, removing the BRCT1 or BRCT2 domains, and introducing phospho-mimetic mutations into the CTR of XRCC4 increased the critical concentration but did not fully prevent LLPS of X4-BRCTs in the presence of XLF (**Fig. S20**). However, XRCC4 alone or in complex with the XID_LIG4_, and the X4liCTR-BRTCs complex did not undergo LLPS with XLF but instead formed filamentous aggregates. Notably, removing both the CTR and the BRCT binding region of XRCC4 (XRCC4_1-151_) suppressed phase separation.

Many transient interactions of the CTRs of XLF and XRCC4 are mediated by electrostatic interactions (**Fig. 2i, 3d**), which are expected to be sensitive to ionic strength. Indeed, increasing the salt concentrations above physiological conditions ([KCl] > 300 mM) effectively dissolved the droplets (**Fig. S17b**). On the other hand, weakening hydrophobic interactions with hexane-1,6-diol only slightly altered the formation of condensates (**Fig. S17c**). Taken together, these data demonstrate that the multivalent interaction network of the CTRs of XLF and XRCC4 is the driving force of the XLF-X4L4 LLPS *in vitro*.

The dramatic concentration increase of specific proteins in condensates may impact the kinetics and the molecular mechanism of NHEJ. Hence, we examined the effect of LLPS on NHEJ ligation activity using a simplified model condensate *in vitro* that comprised either X4L4 alone or in a mixture with XLF (**Fig. 4f-h**). The concentration of the crowding agent (PEG) was increased to induce LLPS, which induced a dramatic stimulation of DNA-end ligation. At the critical concentrations (6% and 4% for X4L4 and X4L4-XLF, respectively), the droplets formed and accelerated intermolecular DNA ligation, most likely through increased local concentration (mass action) as seen in other coacervate droplets (54, 55). As a control, we added increasing amounts of PEG while monitoring DNA ligation by T4 DNA ligase. This led to a small linear increase in ligated DNA products, slightly favoring intramolecular ligation as a result of the excluded volume effect.

### Effects of relevant client proteins on LLPS of XLF and X4L4

Condensates can recruit and concentrate molecules with low valency, which are often referred to as clients (56). The clients are not the main driving force for LLPS but can partition into condensates and alter its properties (57). This aspect is highly relevant for NHEJ as the pathway involves the assembly and sequential action of a variety of proteins (1, 4, 5). To explore the effect of clients on X4L4-XLF condensates, we investigated interactions of XRCC4-XLF condensates with either Ku or the Artemis C-terminal region (Art_CTR_) as models for client proteins. Ku is highly abundant in nuclei, threads on broken DNA ends, and acts as a hub to recruit other NHEJ proteins (58). Ku can form strong to weak interactions with DNA, XLF and the BRCT1 domain of LIG4 (8, 28, 59). Artemis is an endo-and exonuclease that forms a stable complex with DNA-PKcs and is recruited to DSB sites for DNA-end processing (16, 17, 60). In Artemis, the Art_CTR_, which contains 307 residues, interacts with the DBD of LIG4 (61–64). Here, we refer to Ku as a multivalent client, whereas Art_CTR_ is a monovalent client. In co-localization assays, both Ku and Art_CTR_ behaved as client proteins as they effectively partitioned into the X4L4-XLF droplets (**Fig. 5a**_*upper panel*_, **5d**_*upper panel*_). The partition of Ku is DNA- and XLF-interaction dependent (**Fig. 5a**) as the Ku-binding deficient mutation of XLF (F298A) abolished the co-localization of Ku in X4L4-XLF droplets (**Fig. 5a**). On the other hand, Ku enrichment in X4L4-XLF droplets increased significantly in the presence of DNA (**Fig. 5a**). For Art_CTR_, the partition is due to the interaction with DBD_LIG4_ as removing the DBD from LIG4 impaired the recruitment of Art_CTR_ into the droplets (**Fig. 5d**_*lower panel*_). This suggests that X4L4-XLF condensates can selectively recruit relevant factors to DSB sites inside the droplets.

**Figure 5.**
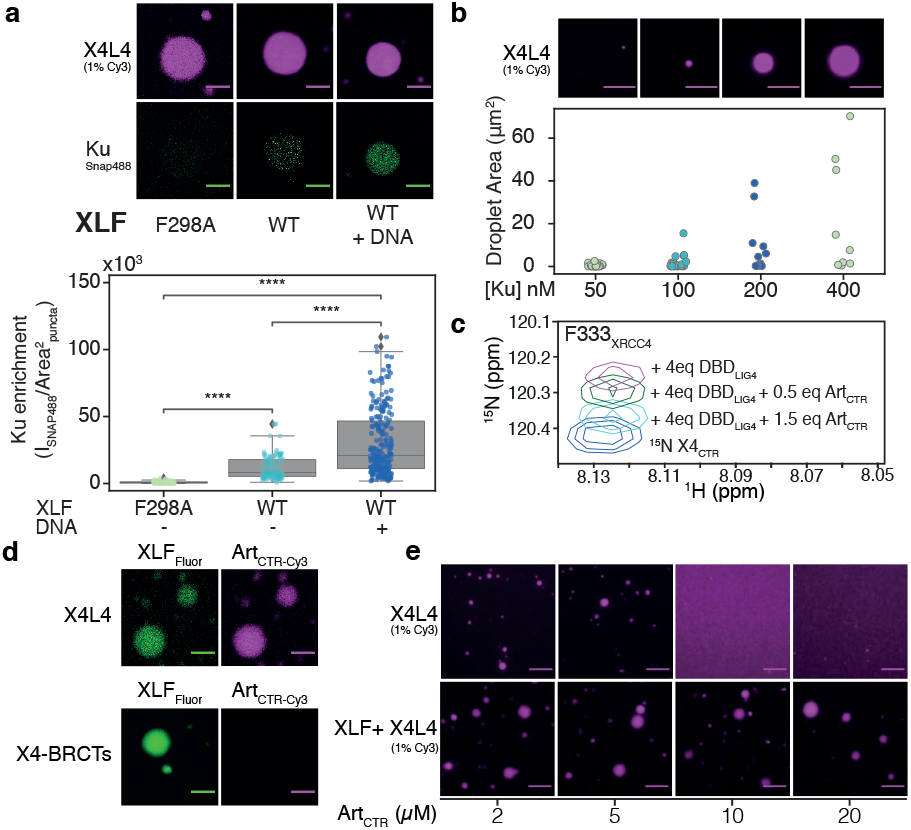
Effect of other NHEJ components on the LLPS of XLF and X4L4. (**a**) Co-localization of X4L4 and Ku in the presence of XLF mutant(F298A, deficient in Ku binding) and WT XLF without or with the addition of DNA (blunt-ended Smal-linearized pUC19). The strip plot and boxplot show the quantification of the Ku enrichment in the puncta across multiple micrographs for each condition. Statistical significance was established using T-test (****P<0.0001). (**b**) *upper panel* : X4L4-XLF droplets with blunt-ended Smal-linearized pUC19 and increasing concentrations of Ku visualized by fluorescence microscope with 1% Cy3-labeled X4L4, *lower panel*: the strip plot of the droplet area across multiple micrographs for each condition, the concentrations of XLF and X4L4 were fixed at 0.5 μM, the experiment was duplicated and analyzed together. (**c**) The inset shows the peak of the F333 residues of XRCC4_CTR_ extracted from the overlaid ^1^H-^15^N HSQC spectra of XRCC4_CTR_ alone or in the presence of DBD_LIG4_ and Art_CTR_. (**d**) Co-localization of Art_CTR_ in X4L4-XLF or XLF-X4-BRCTs droplets. (**e**) Fluorescence microscopy images of X4L4 (1% Cy3 labeled) in the absence or presence of XLF with increasing concentrations of Art_CTR_. The concentrations of X4L4 (complex) and XLF (monomer) were fixed at 5 μM; all scale bars represent 8 μm. Data are represented as mean values and SD.

As expected for a multivalent client (57), the presence of Ku promoted more and bigger X4L4-XLF droplets (**Fig. 5b**). In contrast, adding excess amounts of the Art_CTR_ dissolved the X4L4 droplets (**Fig. 5e**_*upper panel*_), due to competition with the interaction between the CTR of XRCC4 and the DBD of LIG4 at regions around the residue F333 of XRCC4 (**Fig. 3c, 5c**) resulting in valency dilution (57, 65). Yet, the increased number and diversity of transient interactions in X4L4-XLF droplets prevented Art_CTR_ from dissolving the droplets (**Fig. 5e**_*lower panel*_). These results demonstrate that LLPS of X4L4-XLF can be promoted or destabilized by multivalent or monovalent partners, respectively. These results are also compatible with a role of multivalent Ku in the nucleation of X4L4-XLF condensates.

## DISCUSSION

In the past decades, investigations by diverse biophysical approaches have revealed key aspects of the NHEJ mechanism, but the functional roles of IDRs have often been overlooked. Structural biology has often characterized short peptides from IDRs of NHEJ proteins bound to the folded partners with strong-to-intermediate affinities (28, 29, 62, 64), while few studies revealed the functions of IDRs in the context of the full-length proteins (18, 19). Here, we showed that NMR can reveal the dynamics and interactions of the IDRs of core NHEJ factors with atomic resolution in the context of full-length proteins and their complexes. We defined several interactions of XLF and XRCC4 that are driven by their IDRs (**Fig. 6a**). We revealed the motifs in the CTR of XLF that interact with DNA, the folded domain of XLF, and the BRCT1 domain of LIG4. In the case of the CTR of XRCC4, we showed that the CTR can interact transiently with DNA and form a network of electrostatic intra- and inter-molecular interactions. In addition, the XRCC4_CTR_ and Art_CTR_ bind to the DBD domain of LIG4 in a competitive manner. Using cellular models, we also demonstrated that the CTRs of XLF and XRCC4 promote robust cellular NHEJ activity.

**Figure 6.**
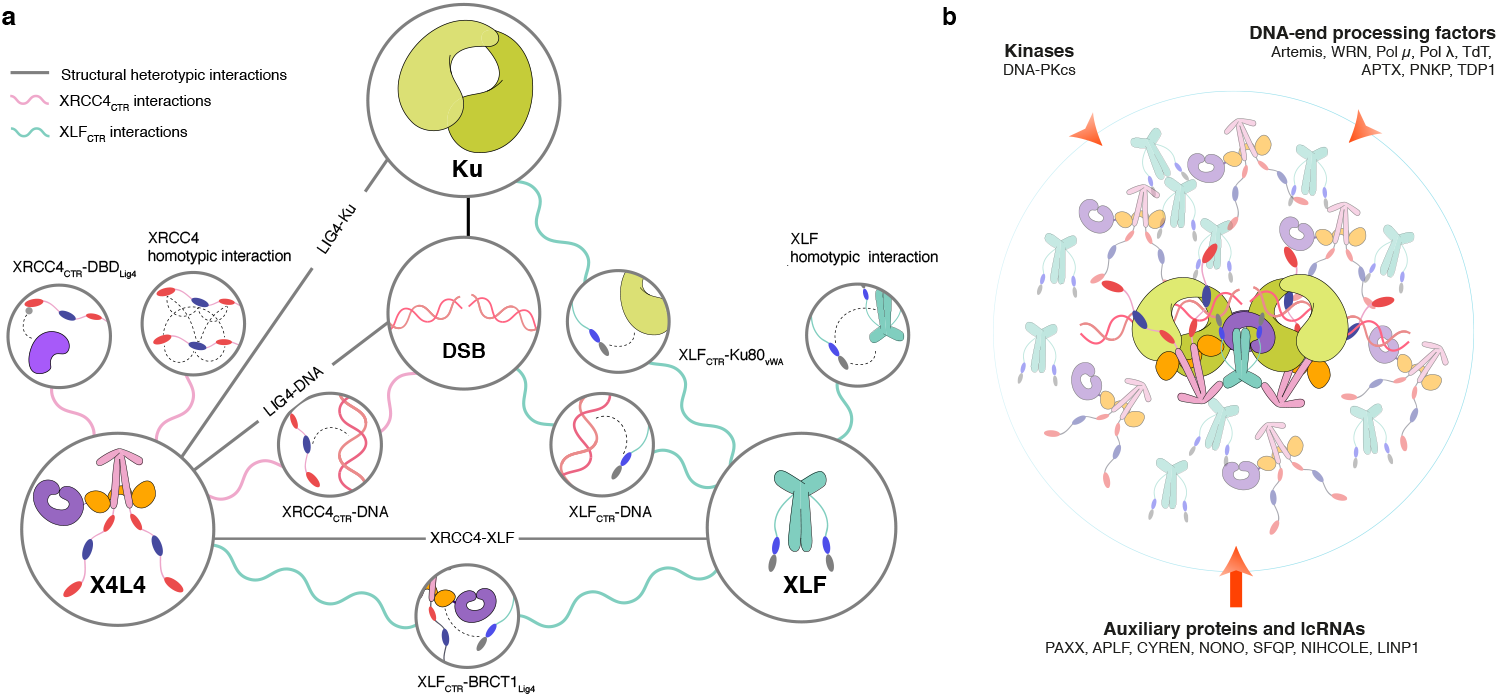
Phase separation from the multivalent interactions of the CTR of XLF and XRCC4 and their role in NHEJ. (**a**) The summary of multivalent interactions of XLF and XRCC4 CTRs characterized in this study (pink and blue wavy lines) as well as previously characterized interactions (straight grey lines). (**b**) Condensates may form and enhance the assembly of NHEJ super-complexes, DNA tethering, rates of enzymatic reactions, and the recruitment of other NHEJ effectors through multivalent interactions of the IDRs, extending the proposed synaptic models obtained from single-particle cryo-EM and single molecule analysis (7, 11). This occurs when the local concentrations of XLF and X4L4 (in shadow) are above a critical concentration. DNA-PKcs: DNA-dependent protein kinase catalytic subunit, WRN: Werner syndrome protein, Pol *μ*: DNA polymerase *μ*, Pol */*: DNA polymerase */*, TdT: terminal deoxynucleotidyl transferase, APTX: aprataxin, PNKP: bifunctional polynucleotide phosphatase/kinase, TDP1: tyrosyl-DNA phosphodiesterase 1, PAXX: paralog of XRCC4 and XLF, APLF: aprataxin and PNK-like factor, CYREN: cell cycle regulator of non-homologous end joining, NONO: non-POU domain-containing octamer-binding protein, SFPQ: splicing factor, proline- and glutamine-rich, lncRNAs: long non-coding RNA (1, 4, 5).

What could be the molecular mechanism underlying the important roles of these CTRs for NHEJ activity? The multiple weak interactions driven by the IDRs can increase local concentrations at DNA repair site through self-enrichment, which would, in turn, increase rates of molecular collisions and favor ligation or act as hubs for the recruitment of accessory factors important for DNA end processing (5, 18, 66). Low-affinity and promiscuous interactions of IDRs can promote rapid but reversible and exchangeable DNA binding, providing dynamic adaptability to the NHEJ process. As IDRs have the ability to move and screen space more freely, they can facilitate and accelerate the assembly of NHEJ super-complexes akin to the fly-casting mechanism (67), which may explain why IDRs accelerate the assembly of NHEJ synaptic complexes (7, 9, 19, 68).

Importantly, we provided evidence that this network of weak multivalent interactions readily and at relatively low concentrations leads to LLPS of X4L4 and XLF *in vitro* (**Fig. 6b**). We showed that these condensates can recruit other NHEJ factors, DNA, and dramatically promote DNA ligation, likely governed by mass action, an effect previously observed in the context of local enrichment of proteins by networks of interactions of IDRs (48, 55). An interesting aspect of protein-DNA co-condensates is that they can have the ability to generate forces that can pull DNA together, which could be highly relevant to the NHEJ mechanism (69).

Can NHEJ components form condensates in response to DSBs in cell nuclei? XLF and X4L4 with endogenous expression levels distribute homogeneously throughout the nucleus (70, 71). However, upon irradiation, XLF and X4L4 are in fast dynamic exchange between DSB sites and the surrounding as their half times of recovery (t_1/2_) in the FRAP experiments are shorter than 10 seconds (70, 71). In addition, numerous XLF and X4L4 clusters can be observed in cells treated with radiomimetic drugs followed by cellular fixation and visualized by super-resolution microscopy (28, 72). Those clusters have been interpreted as XLF and XRCC4 filaments but could instead result from condensates. It is also known that 53BP1, a histone DNA damage-induced mark reader, shares functions with XLF, and forms liquid compartments around sites of DSBs, that facilitate NHEJ (52, 73, 74). Interestingly, two other proteins, SFPQ (Splicing Factor, Proline- and Glutamine-rich) and NONO (Non-POU domain-containing Octamer-binding protein), known to form paraspeckles in the nucleoplasm (75) functionally overlap with XLF in DSB repair by NHEJ (76). Collectively, those results point toward a plausible intervention of condensates in NHEJ at the DSB sites in cells, a hypothesis that requires further investigation.

We dissected the role of individual interactions and found different critical concentrations for phase separation of different constructs and mutants (**Fig. 4e, S19, S20**), which suggests that LLPS of NHEJ could be triggered and/or dissolved by altering the number of multivalent interactions either through post-translational modifications such as sumoylation (23, 77) or recruitment of auxiliary factors such as APLF and long non-coding RNAs (lncRNAs) (53, 78, 79) as seen in other biological systems (80). Indeed, emerging evidence suggests that NIHCOLE or LINP1 lncRNAs stimulate NHEJ through condensate formation (53, 79), yet, the driving forces at the molecular level are unknown. Further, we provided the atomic information about the multivalent interactions of the CTRs of XLF and XRCC4 that will allow the generation of various separation-of-function mutations to test LLPS of NHEJ components directly in cells. Such experiments represent a great challenge due to the high functional redundancy of several NHEJ effectors (73, 81–84), the control of DSB induction (85), and the interplay with other DSB repair pathways such as alternative non-homologous end-joining or homology-driven pathways (2, 86).

In summary, we reveal a multivalent interaction network between NHEJ proteins and nucleic acids mediated by the CTRs of XLF and XRCC4 that are important for robust cellular NHEJ activity as well as for the formation of LLPS *in vitro*. We propose that the local enrichment of NHEJ proteins is a key feature of the mechanism of higher-order synaptic and catalytic assembly for DNA end processing and ligation. Moreover, these hubs or condensates could serve as a flexible platform for the timely recruitment of NHEJ factors and stimulating DNA ligation while protecting the DNA ends from undesired enzymes. The minimal components and molecular interactions underlying NHEJ condensates identified here will serve as a foundation for further *in vitro* and cellular studies. Finally, the emerging understanding of the role of NHEJ disordered protein regions offers an additional dimension in the way NHEJ can be controlled in space and time, and pave the way for potential biomedical applications.

## METHODS

### Plasmid constructs

The WT XLF open reading frame (ORF) was coupled with 6xHistag and TEV-cutting motif at the N-terminus and subcloned into pET16b. XLF mutants were generated by site-directed mutagenesis from the WT XLF. The XLF_CTR_ (229-299) coding sequence was coupled with the SUMO tag and TEV-cutting motif at the N-terminus and subcloned into pET-22b(+). The XRCC4 ORF was coupled with 6xHistag and TEV-cutting motif at the N-terminus and subcloned into pET28b(+). The XRCC4_CTR_ (200-336) coding sequence was coupled with the MBP tag and TEV-cutting motif at the N-terminus and subcloned into pMAL-c5x. The vWA domain of Ku80 (residues 1-245) coding sequence was coupled with MBP tag, 8xHis-tag and TEV-cutting motif at the N-terminus and subcloned into pMAL-c5x. XLF_CTR_, XRCC4_CTR_, MBP-BRCTs cysteine-free and MBP-BRCT1_654-759_ coding sequences were synthesized and sequenced by the Gene Universal Company. Refer to S4 for the comprehensive list of plasmid constructs used in this study.

### Protein expression and purification

#### XLF and XLF mutants

the plasmids containing the XLF and XLF mutants were transformed into *E. Coli* Novagen BL21 Rosetta™(DE3) bacteria. The bacteria were grown at 37 °C in LB containing 75 μg/ml ampicillin and 25 μg/ml chloramphenicol until reaching an optical density at 600 nm between 0.6 and 0.8 and were subsequently induced with 500 μM isopropyl b-D-1-thiogalactopyranoside (IPTG) and incubated overnight at 18 °C. The pellets were collected by centrifugation at 12000×g at 4 °C for 15 minutes and sonicated in buffer B1 (25 mM TrisHCl pH 7.5, 1000 mM NaCl, 2 mM DTT, 1 mM EDTA, 10 mM Imidazole, 1 mM PMSF supplemented with cOmplete™ EDTA-free Protease Inhibitor Cocktail). The supernatant was obtained by centrifuging at 35000×g for 5 minutes at 4 °C and loaded into the prepacked HisTrap™ excel 5ml column preequilibrated with buffer A1 (25 mM TrisHCl pH 7.5, 50 mM NaCl, 2 mM DTT, 1 mM EDTA, 1 mM PMSF) at the rate of 1ml/min and was washed extensively with buffer B1 until obtaining the stable UV signal. In the next step, the column was equilibrated with A1 buffer until the conductivity was smaller than 10 mS/cm. Then, the HiTrapTM 5ml Heparin HP column was attached below the Ni-column. The protein was eluted from Ni-column to Heparin column with 25 ml of buffer A2 (50 mM TrisHCl pH 7.5, 50 mM NaCl, 2 mM DTT, 1 mM EDTA, 300 mM Imidazole, 1 mM PMSF). Then the Ni-column was detached and the inlet pump was changed to A1. The protein was eluted from Heparin column with a buffer gradient (0-60% B1 in 40 mins). The partially truncated of XLF corresponded to the first peak and the second peak was the intact XLF. The His-tag was cut by adding 5% tobacco etch virus protease (TEV) (w/w) into the XLF sample while dialyzing in buffer A1 overnight. The following day, the protein was passed through Ni-column to get rid of uncut XLF and TEV. After that, the follow-through fractions which contained XLF were concentrated and passed through a SEC 26/60 S200 column preequilibrated with NMR buffer (20 mM Bistris pH 6.5, 150 mM KCl, 1 mM DTT, 1 mM EDTA). The fractions corresponding to the peaks were identified by 10% precast polyacrylamide gel, pooled together, and concentrated using 30 kDa MWCO concentrator, flash frozen in liquid nitrogen and stored at −80 °C. The ^15^N and ^15^N/^13^C uniformly labeled samples were prepared using the Marley method (87) with ^15^NH_4_Cl and ^13^C glucose as the sole nutrient sources of nitrogen and carbon, respectively. XLF_CTR_ were expressed and purified using the same protocol as XLF with the exception that the 16/60 S75 column was used in the size-exclusion chromatography.

#### XLF_1-224_ and XLF_1-261_

the plasmids were transformed into *E. coli* Novagen BL21 Rosetta™(DE3) bacteria. XLF_1-224_ and XLF_1-261_ proteins were purified by nickel-immobilized metal affinity chromatography followed by a reversed Ni-column after adding TEV. Subsequently, the proteins were further purified and exchanged into NMR buffer by passing through a S200 26/60 size-exclusion column pre-equilibrated with NMR buffer.

#### XRCC4, XRCC4_1-203_ and XRCC4_1-151_

the plasmids were transformed into *E. Coli* Novagen BL21 Rosetta™(DE3) bacteria. The non-labeled, ^15^N and ^15^N/^13^C uniformly labeled XRCC4 constructs were expressed using the same protocol as XLF. XRCC4, XRCC4_1-203_ were purified using the three-step protocol. Nickel-immobilized metal affinity chromatography followed by Q HP anion exchange chromatography. Next, the samples were buffer exchanged into the NMR buffer. XRCC4_1-151_ was purified using the two-step protocol: nickel-immobilized metal affinity chromatography followed by S200 size-exclusion chromatography in NMR buffer.

#### XRCC4_CTR_

the plasmid was transformed into *E. coli* Novagen BL21 pLysS(DE3) bacteria. The non-labeled, ^15^N and ^15^N/^13^C uniformly labeled XRCC4_CTR_ proteins were expressed using the same protocol as XLF. The XRCC4_CTR_ supernatant was loaded into the prepacked MBPTrap™ HP 5ml column preequilibrated with buffer A1 at the rate of 1ml/min and was washed extensively with buffer B1 until obtaining the stable UV signal. The protein was eluted from the column with buffer 70% A2-MBP (50 mM TrisHCl pH 7.5, 50 mM NaCl, 2 mM DTT, 1 mM EDTA, 30 mM Maltose) + 30% B1. The MBP tag was cut by adding 5% tobacco etch virus protease (TEV) (w/w) into the protein sample while dialyzing in buffer A1 overnight. The following day, the protein was passed through a HiTrap Heparin column to remove MBP and TEV. The fractions corresponding to the XRCC4_CTR_ were concentrated and passed through a SEC 16/60 S75 column preequilibrated with NMR buffer.

#### MBP-BRCT1

the protein was first purified by passing through a HisTrap™ excel 5ml column and then followed by SEC with a 26/60 HiLoad Superdex 200 column to remove misfolded proteins. The MBP and His tags were removed by adding TEV, incubated at 4 °C overnight and passed through the Ni-column. Finally, the protein was loaded into a 16/60 HiLoad Superdex 75 column to exchange into NMR buffer.

#### X4-BRCT1-XID, X4-XID-BRCT2, X4-BRCTs complex

A 1L LB cell pellet of XRCC4 was mixed with a 1L LB cell pellet of different MBP-BRCT constructs. The complexes were purified by a two-step protocol: affinity chromatography by a MBPTrap™ HP 5 ml column, MBP removal by TEV and followed by size-exclusion chromatography. The ^15^N or ^15^N/^13^C uniformly labeled samples were made by mixing a pellet equivalent to 0.5L M9 cell pellet with 1L LB cell pellet of different MBP-BRCT constructs and purified using the same protocol as the non-labeled samples.

#### MBP-Ku80_vWA_

the protein was expressed at 16 °C for two days using the Studier auto-induction protocol (88) and purified using a similar protocol as the MBP-BRCT1 protein.

#### X4L4

the protein was expressed in a similar protocol as MBP-Ku80_vWA_ and purified using 2-step protocols: the Ni-affinity column followed by size-exclusion chromatography in NMR buffer.

#### Art_CTR_

the protein was expressed in a protocol similar to XLF and was captured using a gravity column packed with glutathione sepharose. The protein was released from the column by incubating the resin with PreScission protease overnight, followed by size-exclusion chromatography in NMR buffer.

#### SNAP-Ku (His tag-TEV-Ku80/SNAP-Ku70)

the protein was purified as described for His-tag-TEV-Ku80/Ku70 (28).

### Fluorescence labeling of XLF, X4-BRCTs, XRCC4 and X4L4

The fluorescence dye was incorporated into the protein via maleimide linkage that reacts with cysteine residues. For XLF and X4-BRCTs/X4L4, Fluorescein (Cayman Chemical) and Cy3 (Cytiva) dye was used respectively to react with the solvent-exposed cysteine. For labeling reactions, the proteins were exchanged into buffers containing 50 mM Tris pH 7.5, 150 mM NaCl and 5-fold excessive fluorescence dye were added and incubated overnight at 4 °C. The following day, the mixtures were dialysis excessively 3 times into buffer containing 50 mM TrisHCl 7.5, 150 mM NaCl. The proteins were concentrated, frozen and kept at −80 °C. The 30mer Cy5 labeled DNA was purchased from Eurogentec.

### Microscopy imaging of liquid-liquid phase separation

Unless indicated otherwise, all the LLPS imaging was carried out with buffer including 50 mM TrisHCl 7.5, 150 mM NaCl, 1 mM DTT, 1 mM EDTA and 12.5% Ficoll 400. The components were mixed as indicated concentration and transferred into a cover-glass bottom 384-well black plate (Dutscher) and were then imaged immediately at room temperature using fluorescence microscopy performed on an inverted Zeiss Z1 microscope equipped with a ×60 oil-immersion objective. For the co-localization and FRAP experiments, a Zeiss LSM 710 META laser scanning confocal microscope equipped with an ×63 oil-immersion objective and operated at 20 °C. LSM Zen 2012 Software was used to control and set up experiments. To prevent evaporation and droplet drifting, imaging chambers attached on cover slide (Bio-Labs secureSeal™) were used to contain the samples. Images were analyzed using Fiji (89). The FRAP experiments were performed at 20 °C and the photobleaching of X4L4-XLF_fluorescein_ droplet was achieved using a circular region of interest that covers the whole droplet and a 488 nm laser. Fluorescence recovery was monitored by acquiring images (512×512 pixels) every second at a scanning speed of 0.48 *μ*s per pixel for up to four minutes. The data were analyzed using ImageFRAP (https://imagej.net/Analyze_FRAP_movies_with_a_Jython_script).

### Biochemical assay of DNA-end ligation under LLPS condition

Reaction mixtures (10 *μ*l) contained 40 ng of Xbal linearized (cohesive ends) pUC19 plasmid, 10 mM MgCl_2_, 1 mM ATP, 150 mM NaCl, 50 mM TrisHCl 7.5, 1 mM EDTA, 1 mM DTT, 2 μM X4L4 or 8 U/μL T4 DNA ligase (New England BioLabs) and an increasing amount of PEG 8k (w/v) as indicated. 2 μM XLF were added when indicated. The reactions were incubated at room temperature for 5 minutes. Then, the mixtures were deproteinized by adding pronase E (4 μg/μl) and 1X purple gel loading dye (New England BioLabs) and incubated at 55°C for 30 minutes. The reaction products were analyzed by 0.8% agarose gel electrophoresis using TBE buffer and stained with SYBR Green I. Gel images were visualized and quantified by Gel Doc XR+ and Image Lab™. Each reaction was carried out in triplicate. The reactions were also monitored by a fluorescence microscope with a 60X magnification objective using similar conditions but without adding ATP and MgCl_2_.

### Cellular GFP reporter NHEJ assay

The HEK293 parental, XLF-KO, and XRCC4-KO/XLF-KO cell lines containing the EJ7-GFP reporter were previously described, along with the Cas9/sgRNA plasmids (7a and 7b) used to induce the DSBs in the reporter (23, 90). The XRCC4 and XLF expression constructs were inserted in the pCAGGS-BSKX expression vector, which is also the empty vector (EV) control (91). The reporter cell lines were seeded at 0.5×105 onto a 24-well plate. The cells were transfected on the following day with 200 ng of the 7a and 7b sgRNA/Cas9 plasmids, along with 50ng of the XRCC4 expression vector or EV control, and 50 ng of the XLF expression vector or EV control. Each well contained 1.8 *μ*L of Lipofectamine 2000 (Thermofisher) in 0.5 ml of antibiotic free media. Cells were incubated for 4-hours with the transfection mixture, then washed and fed with complete media. Parallel transfections using a GFP expression vector and EV control (200 ng each) in place of the Cas9/sgRNA plasmids were used to normalize for transfection efficiency. The cells were fixed with 3% formalin 3 days after the transfection and analyzed with the ACEA Quanteon (Agilent NovoExpresss Version 1.5.0), as described (22, 92). For immunoblotting, HEK293 cells were lysed with the ELB (250 mM NaCl, 5 mM EDTA, 50 mM Hepes, 0.1% (v/v) Ipegal, and Roche protease inhibitor) and sonicated with the Qsonica, Q800R. Transfections were scaled to a 6-well plate and HEK293 cells, and used EV to replace the sgRNA/Cas9 plasmids. Blots were probed with antibodies for XRCC4 (CBMAB-X0197-YC, 1:2000), XLF (Antibodies ABIN2857060, 1:2000) and MLH1 (Abcam ab92312, 1:1000). Agilent NovoExpress Version 1.5.0 for the Quanteon was used to capture and analyze the flow cytometry data. Statistical analysis was performed with the Prism Version 8.3.0.

### NMR

#### XLF, XLF_CTR_

Unless specified otherwise, the NMR spectra were acquired at 298 K on a Bruker 18.8 T (800 MHz proton frequency) spectrometer equipped with a room-temperature triple resonance (TXI) probe. The NMR data were processed with NMRPipe (93) and analyzed and visualized using NMRFAM-Sparky (94). The buffer used for the samples was 20 mM BisTris pH 6.5, 150 mM KCl, 1 mM EDTA, 1 mM DTT and 5% D2O. To assign the CTR of XLF, a series of 3D BEST HNCA, HNCACB, and HNCO experiments (95–97) was recorded on a ^13^C, ^15^N uniformlylabeled sample of XLF with a concentration of 250 μM. The resonance assignments of XLF_CTR_ were transferred from the full-length XLF.

#### XRCC4_CTR_, XRCC4, X4-BRCTs

Since the ^1^H-^15^N HSQC spectra of XRCC4_CTR_, XRCC4, X4-BRCTs are very similar, the resonance assignment was done on XRCC4_CTR_ and was subsequently transferred unambiguously to the other constructs. A series of 3D BEST HNCA, HNcoCA, HNCACB, HNcoCACB and a series of 3D BEST-TROSY HNcoCA, 3D BEST-TROSY HNcaNh, HNcacoNh experiments were acquired for this purpose (95–97). Moreover, a series of 3D BEST HNCACB, HNcoCACB experiments was acquired on the ^13^C/^15^N uniformly labeled X4-BRCTs sample to confirm the transferred assignment from XRCC4_CTR_ and to determine the chemical shift of the CA and CB resonances. The BRCT1 domain was assigned using the similar series of 3D experiments as for XRCC4_CTR_.

#### NMR relaxation measurement

Unless specified otherwise, the NMR relaxation experiments were acquired on an 800 MHz spectrometer at 298 K using uniformly ^15^N labeled samples. The ^15^N longitudinal relaxation rates (R_1_) were acquired by sampling the decay of the magnetization at 6 different delays (for XLF) or 10 delays (for XRCC4 constructs) between 0 and 1.8 s. The ^15^N transverse relaxation rates (R_2_) were recorded using two different methods: R_2_^echo^ using a single echo while decoupling the protons during the entire relaxation period or R_2_^CPMG^ using Carr-Purcell-Meiboom-Gill (CPMG) train of ^15^N p-pulses interleaved with ^1^H p-pulses (98, 99). Six delays between 4 and 100 milliseconds were used to determine R_2_^CPMG^, while seven delays between 9 and 288 milliseconds were used for the R_2_^echo^ experiments. ^1^H-^15^N heteronuclear (hNOE) were measured by detecting the ^15^N steady-state polarization while saturating the protons with a train of p-pulses and comparing the peak intensities to the ones of an experiment where the magnetization of ^15^N has returned to equilibrium (100).

#### NMR titration experiments

For the DNA-protein titration experiments, DNAs (purchased from Eurogentec) were added to the isotopically labeled proteins after exchanging them into the NMR buffer. For protein-protein titration experiments, the unlabeled proteins were added to the solution of the isotopically labeled proteins. Chemical shift perturbation (CSP) values were calculated for all residues by subtracting the shifts of the proteins in the mixtures with DNA or partner proteins from the shifts of the protein alone, using the following equation: 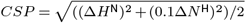

#### Paramagnetic relaxation enhancement

Cysteine-free XLF, XRCC4, and XRCC4_CTR_ plasmids were produced and a single cysteine residue was then introduced at the desired position by site-directed mutagenesis. The proteins were expressed and purified using a similar protocol as the ^15^N labeled proteins. In the final gel filtration chromatography step, a buffer of 25 mM TrisHCl 7.5 and 150 mM NaCl was used. The proteins were concentrated and an amount of 10-fold excess of MTSL (Cayman Chemical) was added. The mixtures were incubated overnight at 4 °C. The following day, the mixtures were dialyzed 3 times into PRE buffer (20 mM Bistris 6.5, 150 mM KCl) to remove the unreactive MTSL completely. The proteins were concentrated, frozen, and kept at −80 °C. The concentration of proteins for the PRE measurements was 50 μM. ^1^H-^15^N HSQC spectra of the proteins with the MTSL probe in the paramagnetic or the diamagnetic state were recorded with 256 complex points in the indirect dimension. A delay of five seconds was used between scans to ensure the complete magnetization recovery. The diamagnetic state of MTSL was generated by adding 10 molecular equivalents of ascorbic acid and incubating overnight. The spectra of the diamagnetic states were recorded the following day. The experiments were carried out at 298 K, on an 800 MHz spectrometer (for XLF and X4-BRCTs) or 600 MHz spectrometer (for XRCC4_CTR_).

### EPR

Cysteine variants of XRCC4 (C218, C299, C337) alone or in complex with BRCTs have been individually incubated with a 10-fold molar excess of DTT. The reducing agents have been removed from each sample with PD-10 desalting columns using 20 mM Bis-Tris (pH 6.5) with 150 mM KCl as the eluting solution. Successively, a 10-fold molar excess of MTSL spin label was added to the protein solutions and the reactions were kept at 4 °C under stirring. After 2 hours, a second addition of the spin label was carried out and the reaction was maintained in the same conditions during 2 additional hours. The unreacted fraction of MTSL was removed from each protein sample using PD-10 desalting columns with the buffer solution reported above. Spin labeled XRCC4 variants were concerted using AMICON centrifugal filters (MWCO = 10kDa) and the protein concentrations were calculated from the UV/vis spectra (A280, ε = 25440 M^-1^.cm^-1^). The spin concentration of each sample has been calculated from the double integral of the EPR spectrum and the relative intensity has been compared to a standard. The labeling yields were in the 80-90% range for all MTSL-labeled XRCC4 mutants. The protein solutions have been successively stored at −80 °C. The XRCC4-BRCTs complexes were labeled in a similar manner. CW-EPR spectra have been recorded at room temperature on a Bruker Elexsys E500 spectrometer equipped with a Super High Q resonator operating at X-band (9.9 GHz). Samples (50μL) have been loaded into quartz capillaries with 80-100 μM of final concentration. The following parameters have been used: microwave power 20 mW; magnetic field modulation amplitude 1G; modulation frequency 100 kHz; field sweep 15 mT; accumulations 4. Simulations of the EPR spectra (**Fig. S21** and **Table S1**,**S2**,**S3**) have been carried out using Simlabel (101) (a GUI of Easyspin (102))

### Small-angle X-ray scattering experiments

The SAXS data of XLF and XRCC4-BRCTs complex were collected at the Swing beamline (Proposal No 20200489) of SOLEIL synchrotron using the size-exclusion chromatography coupled with small angle X-ray scattering (SEC-SAXS). Peak picking and buffer subtraction were done using Chromix in ATSAS-3.0.3-1 package (103). For XRCC4_CTR_, the SAXS data were collected at DIAMOND beamline 21 using the batch method. The buffer was similar to NMR buffer plus 2% glycerol. The dimensionless Kratky plots were analyzed and generated using BioXTAS RAW package (104).

### ASTEROIDS ensemble analysis of the disordered of XRCC4_CTR_

The ensemble of states sampled by isolated XRCC4_CTR_ in solution were characterized using the ensemble selection algorithm ASTEROIDS (47). Initially, ensembles were selected on the basis of experimental heteronuclear backbone chemical shifts (^13^CO, ^13^CA,^13^CB, 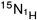,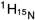). 10000 conformers were initially calculated using a residue-specific statistical coil model of backbone dihedral angle sampling (45, 46). Sidechains were added using the sccomp algorithm (105) and chemical shifts were calculated using Sparta (106). 5 ensembles of 200 structures were selected to match experimental chemical shifts (47), iteratively doping the statistical coil database with dihedral sampling derived from the previous step to aid convergence, as described elsewhere. The resulting sequence-specific chemical shift sampling was then used to calculate 20000 conformers, from which chemical shifts, paramagnetic relaxation (107) and small angle scattering curves (108) were calculated and appropriately averaged over the ensemble as previously described. Finally, representative ensembles of 200 structures were selected to match all experimental data within experimental uncertainty. The contact map is calculated as the log10 of the ratio of the average distance calculated over the selected ensemble and the reference ensemble from the initial pool of structures (107).

## ACKNOWLEDGEMENTS

We acknowledge SOLEIL for provision of synchrotron radiation facilities and we thank Aurelien Thureau for assistance in using beamline SWING. We thank Katrine Bugge, Birthe B. Kragelund and Johan G. Olsen for accessing DL-SAXS facility at Diamond Light Source (grant no. EP*/*R042683*/*1), Nicola Salvi and Nicolas Bolik-Coulon for fruitful discussions as well as Katheryn Meek and Eli Rothenberg for critical reading of the manuscript.

## Funding

This work was funded by the French ANR (ANR-18-CE29-0003 NANO-DISPRO, ANR-17-CE2-0020 NHEJLIG4) and French Ligue against Cancer (équipe labellisée). This work has been supported by the Fondation ARC pour la recherche sur le cancer (grant N°ARCDOC42021120004347), ERC Grant agreement 279519 (2F4BIODYN) to F.F. and 835161 (DynamicAssemblies) to M.B. as well as the Equipex contract ANR-10-EQPX-09 (Paris en resonance). A.K.A was supported by grant R35-GM131780 from the National Institutes of Health. Financial support from the IR INFRANALYTICS FR2054 CNRS for conducting the research is gratefully acknowledged.

## Authors contributions

F.F., M.M., V.B. and M.Bl. conceived the project and obtained the main funding. D.D.V. prepared reagents and performed the NMR and *in vitro* experiments with the help of P.P., Z.W., L.C., G.B. and Z.G.; M.Br., M.M., M.C.-A., J.M.S prepared reagents and performed the cellular experiments. A.B. and V.B. conducted the EPR experiments. M.Bl., J.M.S., Z.G., P.C., P.P., L.C., G.B. and A.K.A. assisted in conceptualization and analysis of data. D.D.V., M.M. and F.F. wrote the manuscript and all the co-authors edited and approved the final version of the manuscript.

## Competing interests

The authors declare no competing interests.

## Data availability

The NMR assignments of the CTR of XLF, XRCC4 as well as BRCT1 domain will be deposited to BMRB (code to be determined). The datasets generated during and/or analyzed during the current study are available from the corresponding authors on reasonable request.

## Supplementary Materials

Supplementary figures: Fig. S1-S21

Supplementary tables: Table S1-S4

## Notes

### Competing Interest Statement

The authors have declared no competing interest.

