## Supporting Information for "Multivalent interactions of the disordered regions of XLF and XRCC4 foster robust cellular NHEJ and drive the formation of ligation-boosting condensates *in vitro*"

### Supplementary Figures

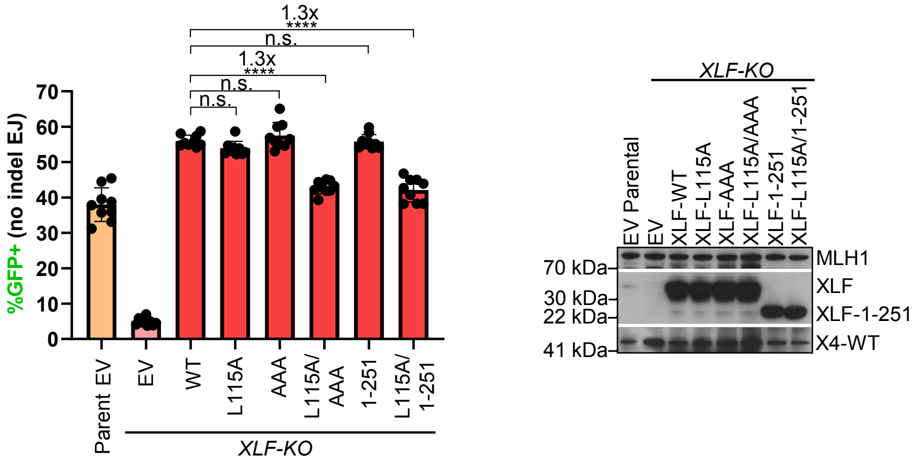


**Figure S1. Effects of XLF mutations on No Indel EJ with endogenous XRCC4.** GFP frequencies are normalized to parallel GFP transfections to account for transfection efficiency. n = 9 biologically independent transfections. Statistics with unpaired two-tailed t-test with Holm-Sidak correction. ****P<0.0001, n.s. = not significant, #x represents fold effect. Immunoblots show levels of XRCC4, XLF-WT, and XLF mutants.

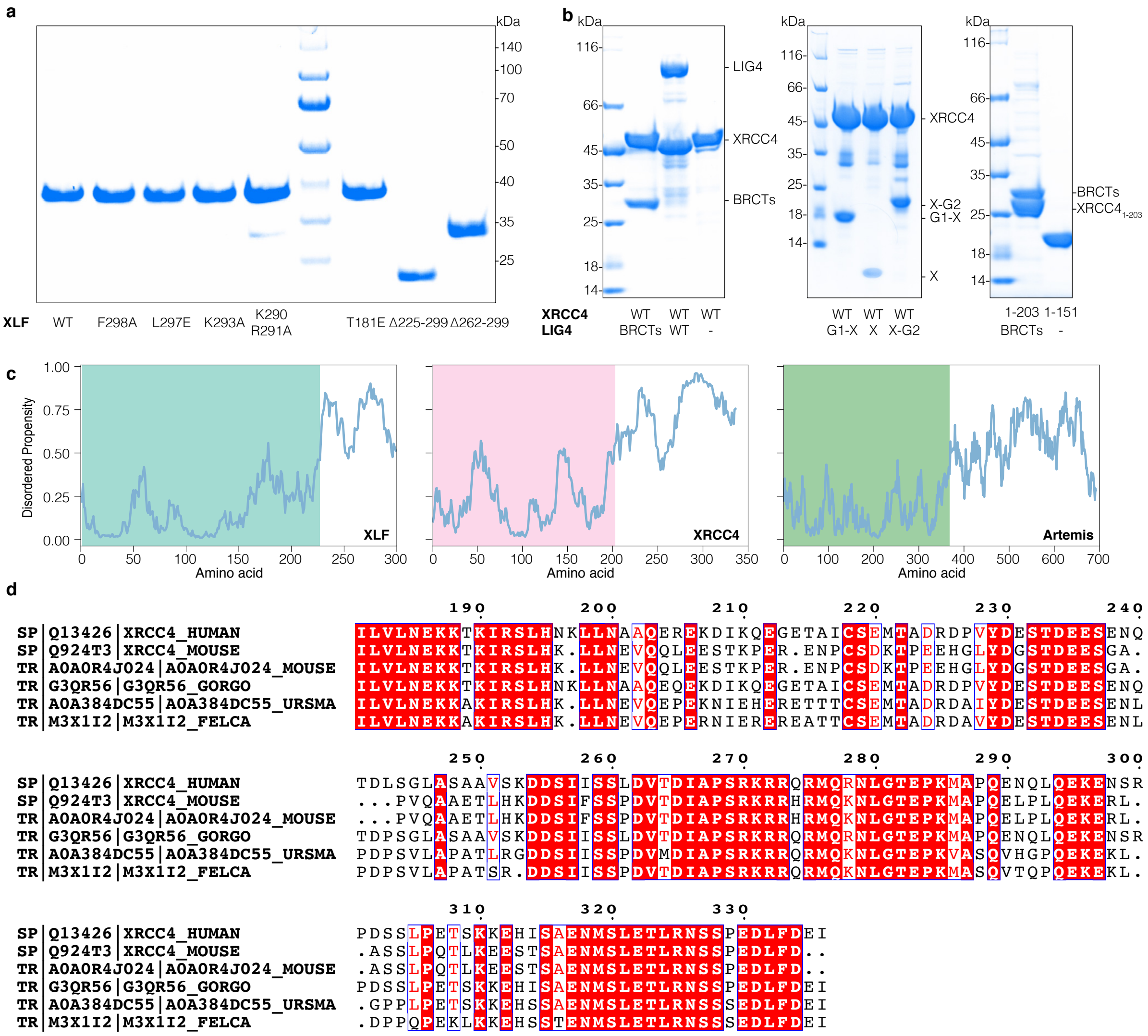


**Figure S2. Protein constructs used in the study.** 10% SDS-PAGE and InstantBlue™ Coomassie Stain analysis of purified proteins: (**a**) XLF wild-type, mutants and truncated constructs, (**b**) XRCC4 wild-type and truncated constructs alone or in complex with different BRCTs domains of LIG4. (**c**) Disordered propensity prediction of XLF, XRCC4, and Artemis using IUPRED2A (1), the regions where the structures are available are shaded in colors. (**d**) Multiple sequence alignment of XRCC4CTR from different species. WT: wild-type, BRCTs: tandem BRCTs domain of LIG4 (residues 654-911), G1-X: first BRCT domain and XRCC4-binding domain of LIG4 (residues 654-802), X: XRCC4-binding domain (residues 739-802), X-G2: XRCC4-binding domain and second BRCT domain of LIG4 (739-911).

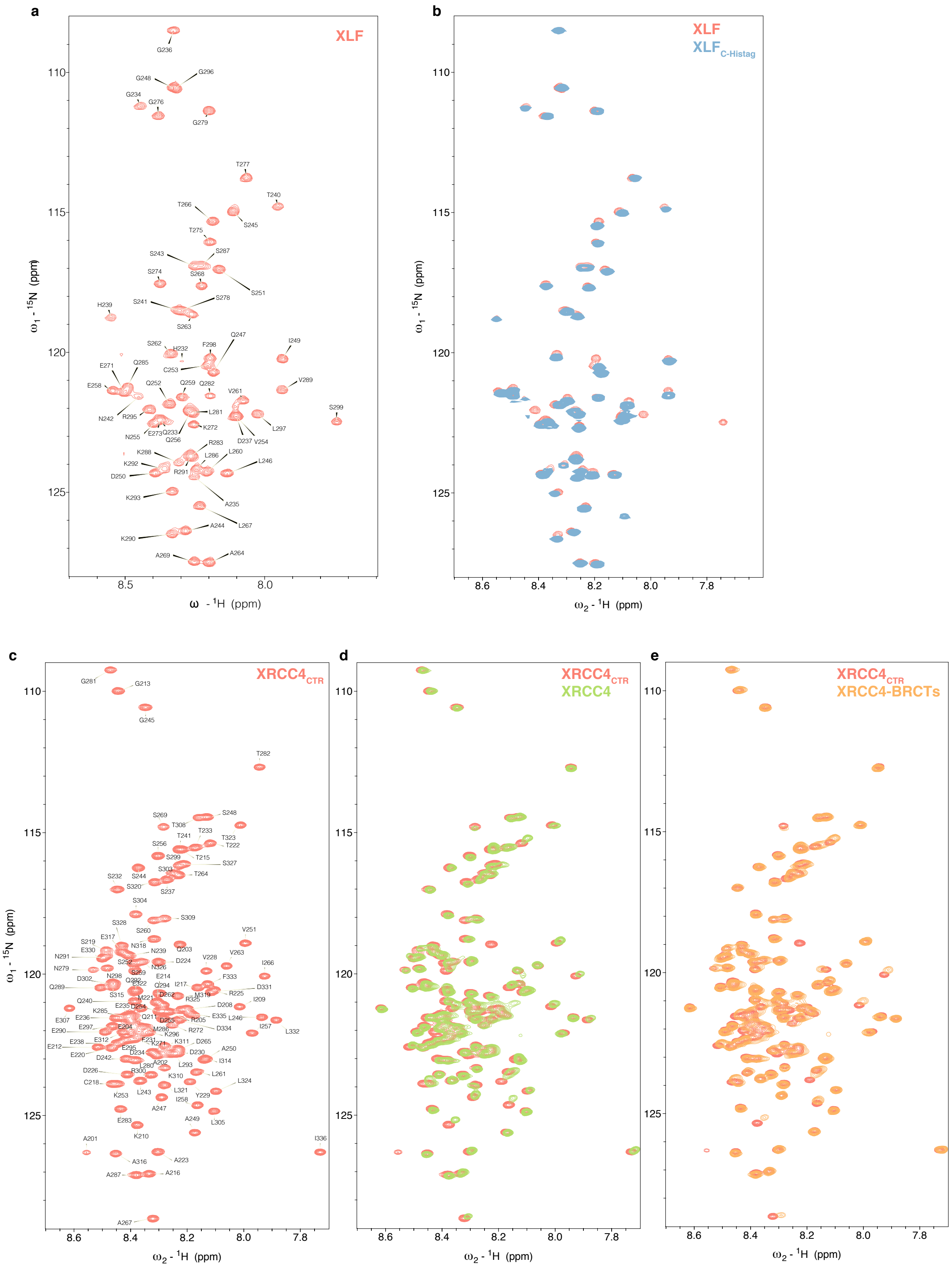


**Figure S3. NMR resonance assignment of XLF and $\mathrm{XRCC4_{CTR}}$.** (**a**) NMR resonance assignment of $^{1}$H-$^{15}$N HSQC spectrum of XLF. (**b**) Overlay $^{1}$H-$^{15}$N HSQC spectra of XLF (red) and XLF with a His-tag at the C-terminus ($\mathrm{XLF_{C\text{-}Histag}}$, blue). (**c**) NMR resonance assignment of $^{1}$H-$^{15}$N HSQC spectra of $\mathrm{XRCC4_{CTR}}$. Overlay $^{1}$H-$^{15}$N HSQC spectra of $\mathrm{XRCC4_{CTR}}$ (red) and XRCC4 (200 µM, monomer, green) (**d**) and XRCC4-BRTCs (70 µM, complex, orange).

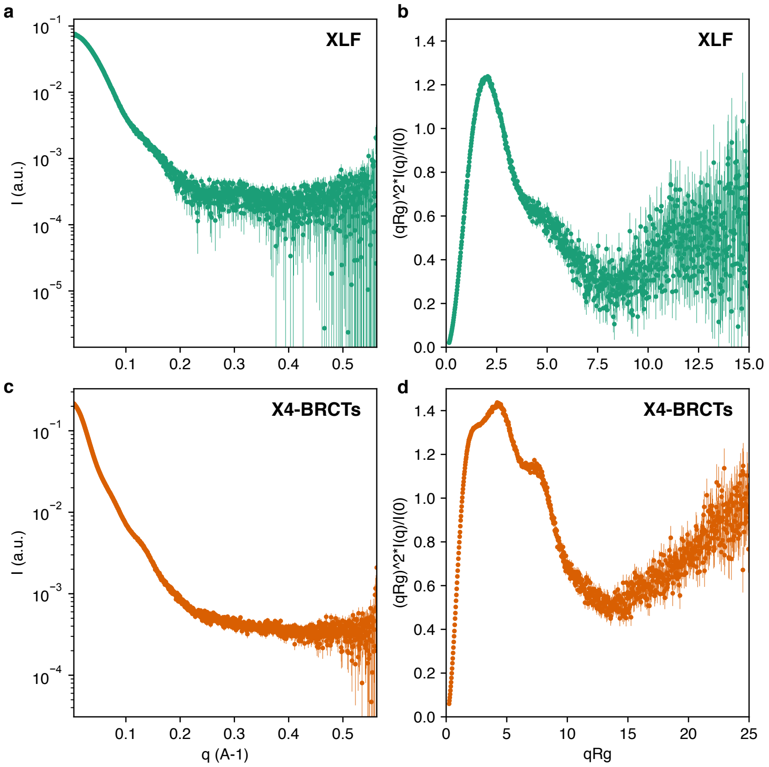


**Figure S4. SEC-SAXS experiments of XLF and XRCC4 in complex with BRCTs domain of LIG4 (X4-BRCTs)** (**a,c**) I(q) in the log scale versus q in the linear scale of XLF and X4-BRCTs respectively. (**b,d**) Dimensionless Kratky plots of XLF and X4-BRCTs, respectively.

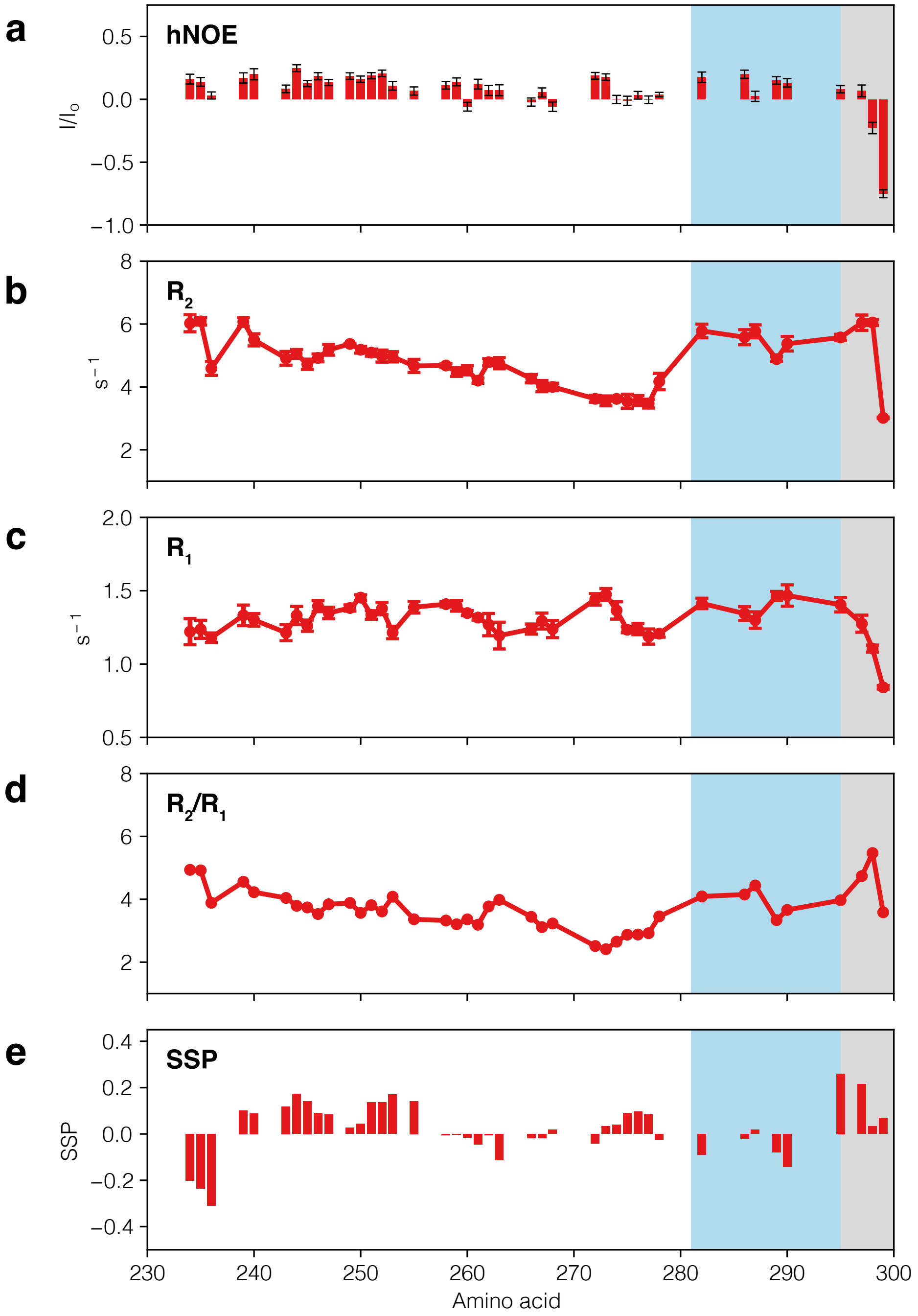


**Figure S5. NMR parameters of XLF.** Only the peaks belonging to the residues in the CTR are visible and analyzed. (**a**) Heteronuclear [$^{1}$H]-$^{15}$N Overhauser effects (hNOE) of XLF. (**b**) CPMG transverse relaxation rate ($R_2$), (**c**) longitudinal relaxation rate ($R_1$), (**d**) $R_2/R_1$ ratio of XLF and (**e**) secondary structure propensity of XLF calculated using Cα and Cβ chemical shift. All parameters were measured on an 800 MHz spectrometer at 298 K; the residues in the DNA-binding motif are highlighted in blue; the residues in the Ku-binding motif are highlighted in gray.

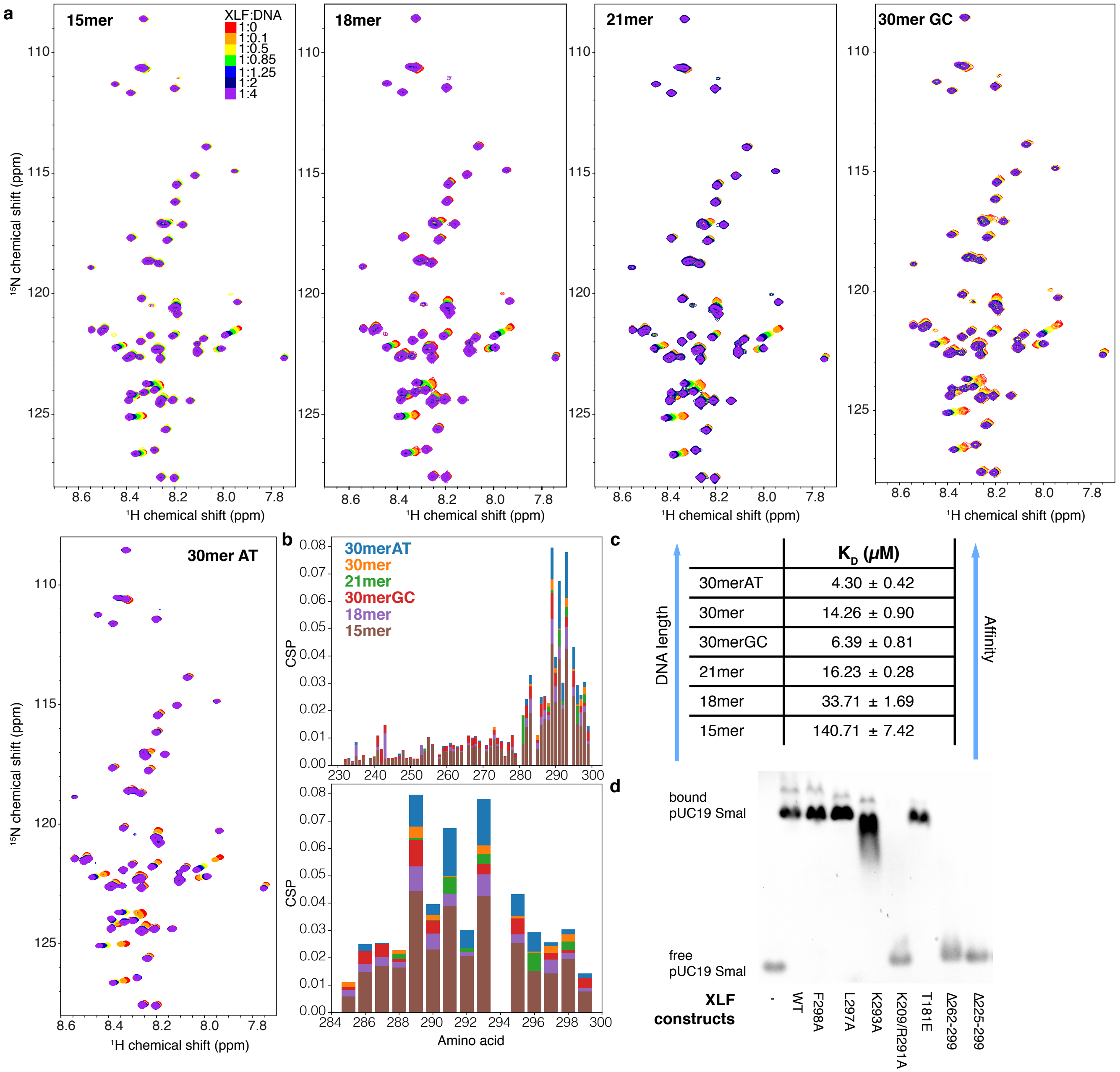


**Figure S6. Titration of XLF with DNA monitored by NMR.** Only the peaks belonging to the residues in the CTR are visible and analyzed. (**a**) Overlay $^{1}H$-$^{15}N$ HSQC spectra of XLF (red) after adding 0.1, 0.5, 0.85, 1.25, 2, and 4 equivalents of five types of DNA as indicated. (**b**) Top: Chemical shift perturbation of XLF before and after adding 4 equivalents of different types of DNA; bottom: as in the top figure but the binding site is enlarged. (**c**) The apparent affinity of XLF ($K_D^{app}$) for DNA by fitting 2D line shape of affected peaks by TITAN software. (**d**) Electrophoretic mobility shift assay (EMSA) of different XLF constructs with DNA (SmaI-linearized pUC19).

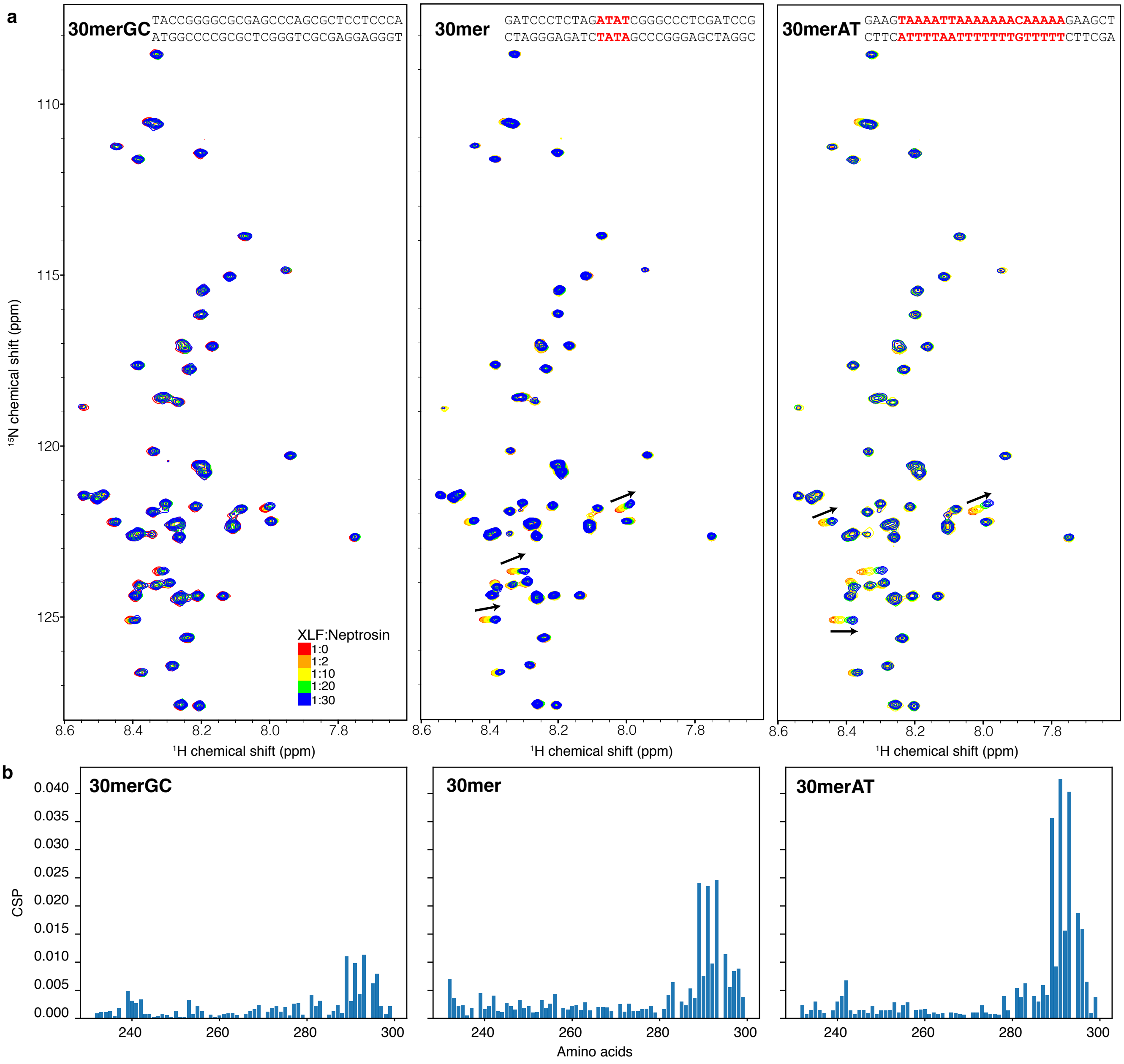


**Figure S7. Netropsin-XLF competitive NMR titration experiments.** Only the peaks belonging to the residues in the CTR are visible and analyzed. (**a**) Overlay $^{1}H$-$^{15}N$ HSQC spectra of XLF in the presence of 4 equivalents (red) of 30mer DNA (GC rich, random, AT-rich) and after adding 2 (orange), 10 (yellow), 20 (green) and 30 (blue) equivalents of Neptrosin, the sequences of DNA are included with the spectra in which the AT-rich regions where Netropsin can bind are highlighted in red. (**b**) Chemical shift perturbation of XLF in the presence of 4 equivalents of three different kinds of DNA before and after adding 30 equivalents of Netropsin.

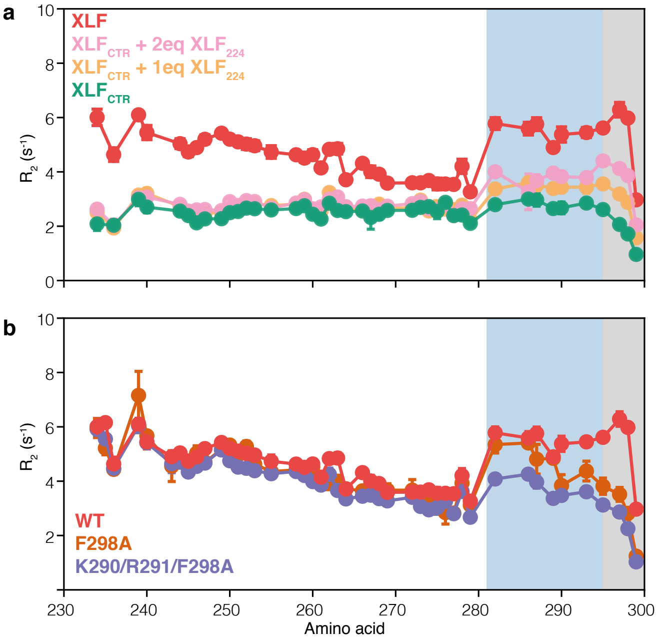


**Figure S8. $XLF_{CTR}$ interacts with the folded domain.** Only the peaks belonging to the residues in the CTR are visible and analyzed. (**a**) Traverse relaxation rate (R2) of $XLF_{CTR}$ after adding 1 (orange) or 2 (violet) equivalents of $XLF_{1\text{-}224}$ compared with full-length XLF (red). (**b**) $R_2$ of XLF in comparison with two other mutants. All the rates were measured on an 800 MHz spectrometer at 298 K, the residues in the DNA-binding motif are highlighted in blue, and the residues in the Ku-binding motif are highlighted in gray.

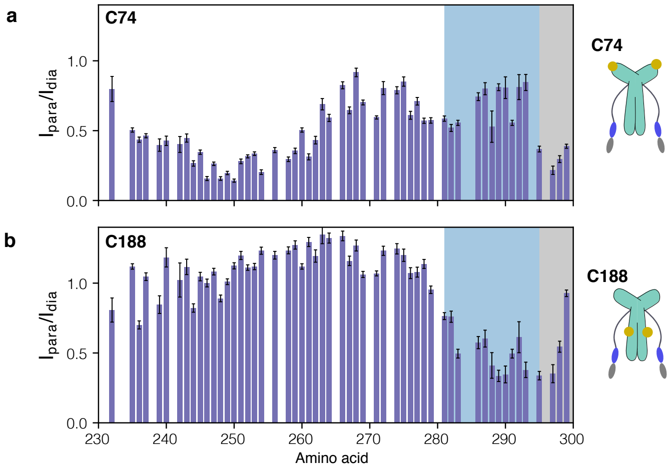


**Figure S9. Paramagnetic relaxation enhancement (PRE) experiments of XLF.** Only the peaks belonging to the residues in the CTR are visible and analyzed. (**a-b**) Peak intensity ratio derived from $^{1}H$-$^{15}N$ HSQC spectrum between the paramagnetic and diamagnetic state of MTSL probe at the positions C74 and C188, respectively. The structural presentation of XLF with the yellow spheres are depicted as MTSL nitroxide at the respective cysteine residues. The PREs were measured on an 800 MHz spectrometer at 298 K, the residues in the DNA-binding motif are highlighted in blue, the residues in the Ku-binding motif are highlighted in gray.

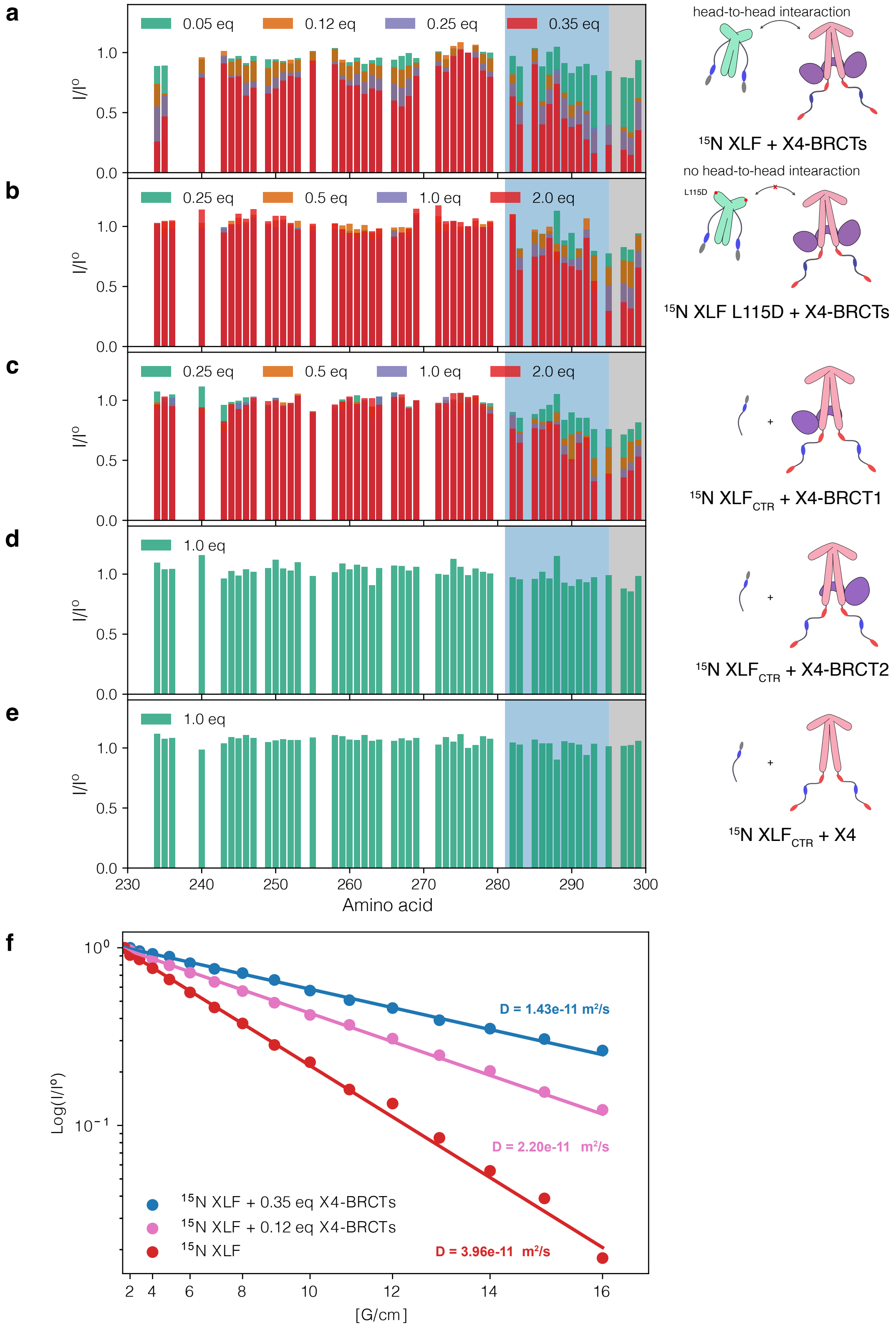


**Figure S10. $XLF_{CTR}$ interacts with the BRCT1 domain of LIG4.** Only the peaks belonging to the residues in the CTR are visible and analyzed. Normalized peak intensity ratios extracted from $^{1}$H-$^{15}$N HSQC spectra between free $^{15}$N XLF (**a**), XLF L115D (**b**), $XLF_{CTR}$ (**c-e**) ($I^{o}$) in the presence of different equivalents of X4-BRCTs (**a-b**), X4-BRCT1 (**c**), X4-BRCT2 (**d**), XRCC4 (**e**) (I). (**f**) Measurement of the diffusion coefficient of XLF using 1D $^{15}$N-edited pulsed field gradient diffusion experiments (2) recorded with 0 (red), 0.12 (pink), and 0.35 (blue) equivalent of X4-BRCTs.

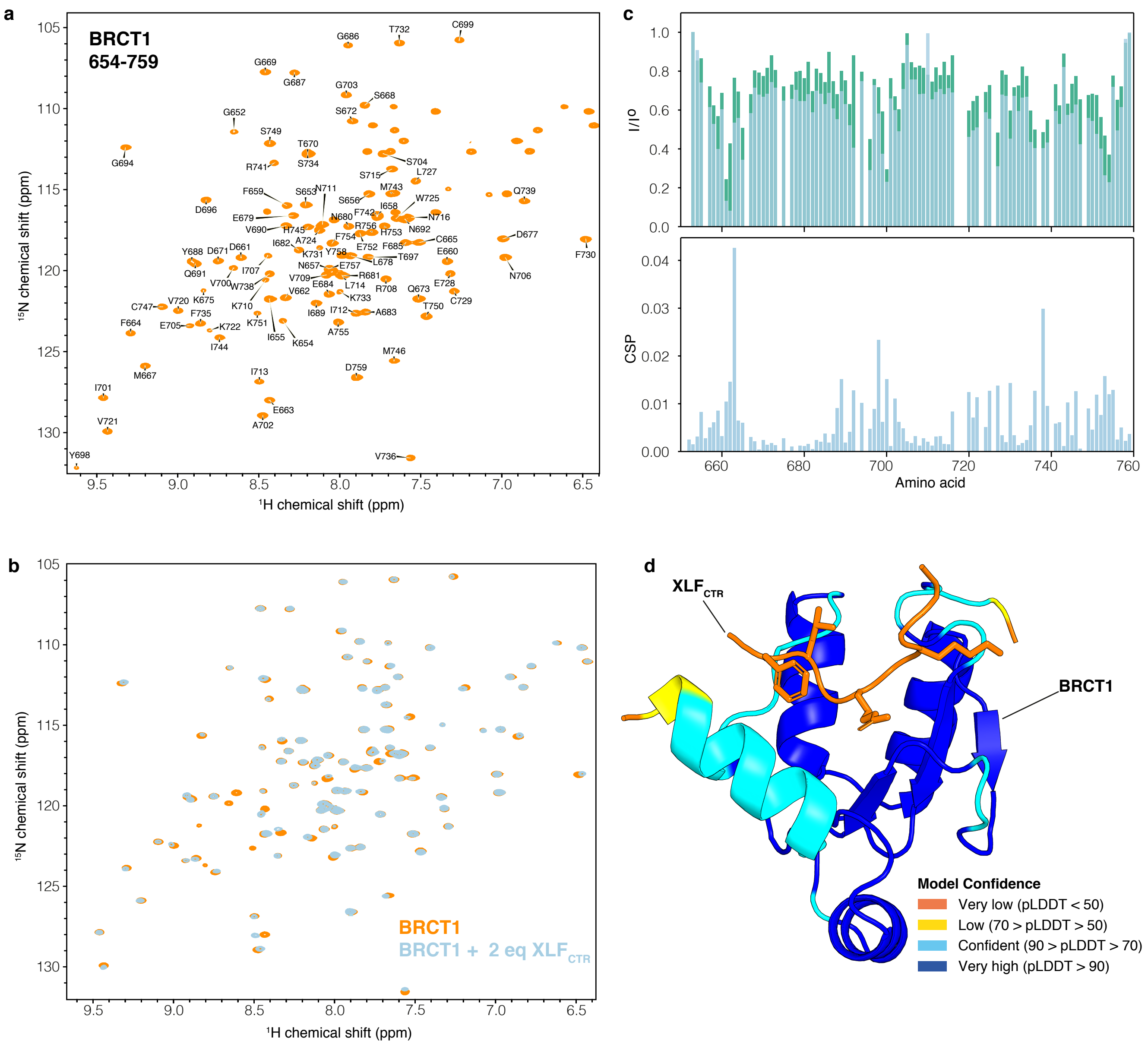


**Figure S11. Identifying the region of BRCT1 that interact with $XLF_{CTR}$.** (**a**) $^{1}$H-$^{15}$N HSQC spectra of BRCT1 (residues 654-759) with resonance assignment. (**b**) Overlay $^{1}$H-$^{15}$N HSQC spectra of BRCT1 in the absence (orange) and in the presence (cyan) of two equivalents of $XLF_{CTR}$. (**c**) *top*: normalized peak intensity ratios extracted from $^{1}$H-$^{15}$N HSQC spectra of $^{15}$N BRCT spectra before ($I^{o}$) and after adding 1 (green) or 2 (cyan) equivalents of $XLF_{CTR}$ (I), *bottom*: chemical shift perturbation (CSP) of BRCT1 after adding 2 equivalents of $XLF_{CTR}$. (**d**) BRCT1-$XLF_{CTR}$ complex predicted using AlphaFold-Multimer (3) implemented on Colabfold (4) with default parameters.

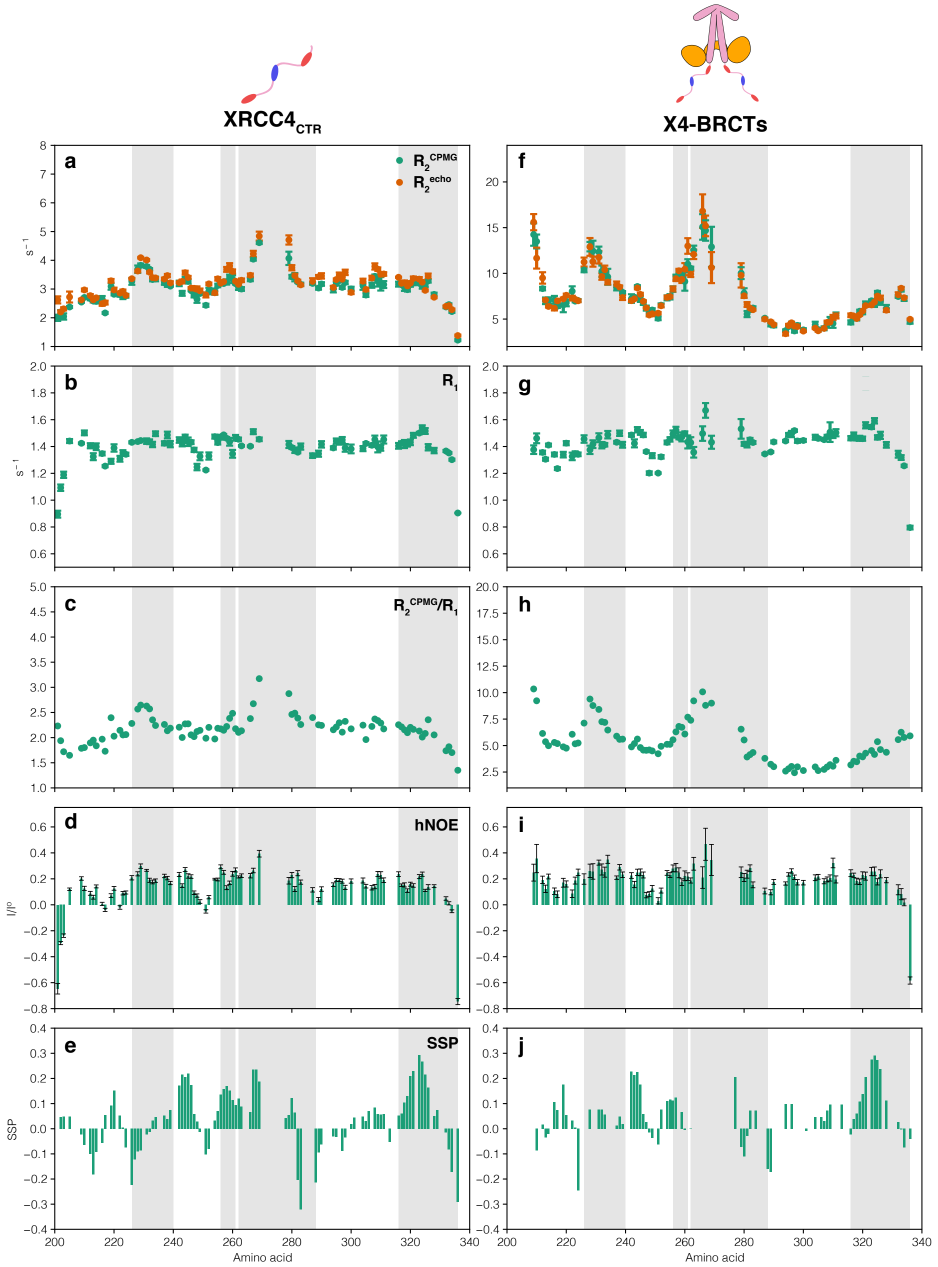


**Figure S12. NMR parameters of $XRCC4_{CTR}$ and XRCC4 in complex with BRCTs.** Only the peaks belonging to the residues in the CTR are visible and analyzed. (**a, f**) Transverse relaxation rate measured under a CPMG train ($R_2^{CPMG}$, in green) or simple echo ($R_2^{echo}$, in orange), (**b, g**) longitudinal relaxation rate ($R_1$), (**c, h**) $R_2/R_1$ ratio, (**d, i**) heteronuclear $[^1H]$-$^{15}N$ Overhauser effects (hNOE), (**e, j**) secondary structure propensity calculated using Cα and Cβ chemical shift of $XRCC4_{CTR}$ and X4-BRCTs respectively. All the parameters were measured on an 800 MHz spectrometer at 298 K, the residues are highlighted in gray as follows: Forkhead-associated-domain Binding Motif (FBM, residues 226-240) SUMO Interaction Motif (SIM, residues 256-261), DNA-PKcs Interaction Motif (DIM residues 262-288), Negatively Charged C-terminal region (NCT, residues 316-336).

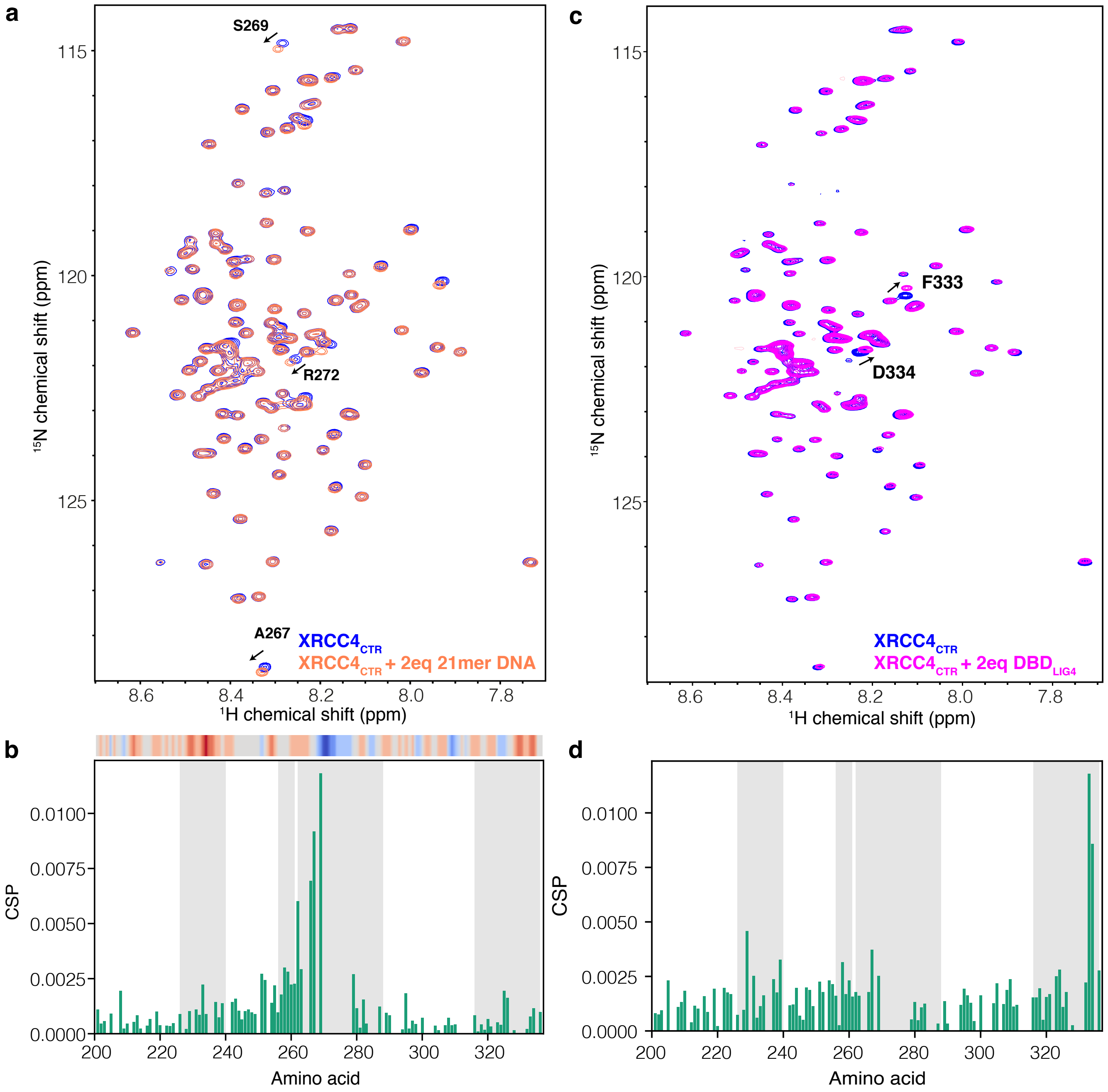


**Figure S13. The CTR of XRCC4 can bind transiently to DNA and the DBD domain of LIG4.** (**a**) Overlay of $^{1}$H-$^{15}$N HSQC spectra of XRCC4$_{CTR}$ alone (blue) and in the presence of 2 equivalents of 21mer DNA (salmon color). (**b**) Chemical shift perturbation (CSP) extracted from the spectra in (**a**), the charge distribution was shown on top. (**c**) Overlay of $^{1}$H-$^{15}$N HSQC spectra of XRCC4$_{CTR}$ alone (cyan) and in the presence of 2 equivalents of the DBD of LIG4 (purple). (**d**) Chemical shift perturbation (CSP) extracted from the spectra in (**c**). The residues are highlighted as in S12.

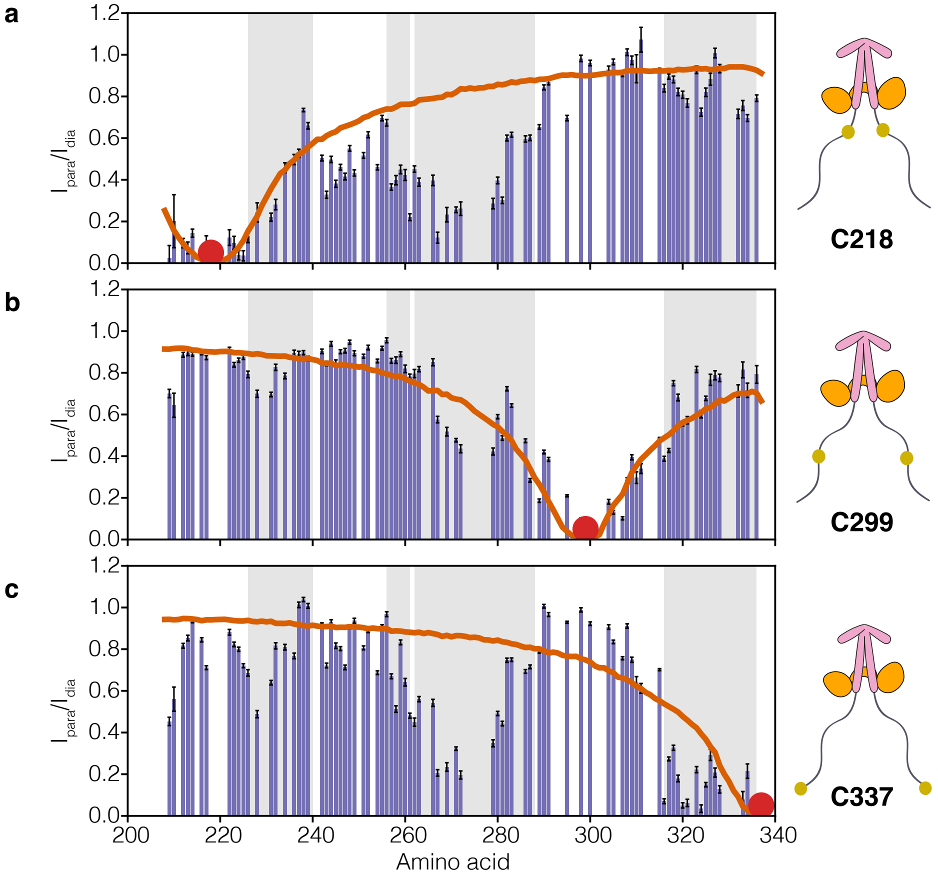


**Figure S14. PRE experiments of XRCC4 in complex with BRCT domains.** Only the peaks belonging to the residues in the CTR are visible and analyzed. Peak intensity ratio derived from $^{1}H$-$^{15}N$ HSQC spectra between paramagnetic and diamagnetic states of MTSL probed at three positions: C218 (**a**), C299 (**b**), C337 (**c**); the red line is the PRE effect calculated from 100 000 random XRCC4$_{CTR}$ conformations using Flexible Meccano (5). The residues are highlighted as in S12.

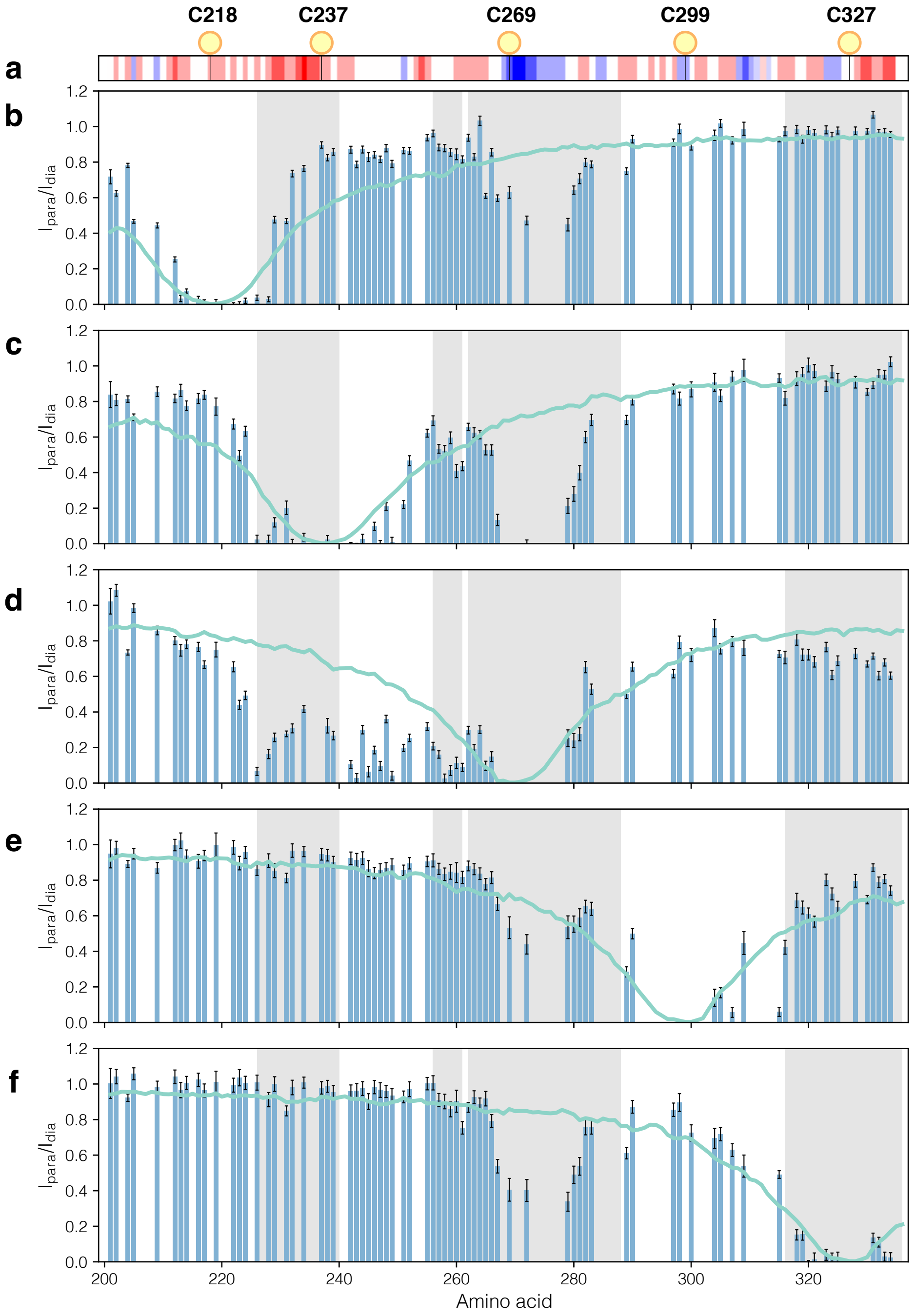


**Figure S15. PRE experiments of $XRCC4_{CTR}$.** (**a**) Charge distribution of $XRCC4_{CTR}$ (three-residue window) with the yellow spheres are depicted as MTSL nitroxide at the respective cysteine residues. Peak intensity ratio derived from $^{1}H$-$^{15}N$ HSQC spectra between paramagnetic and diamagnetic states of MTSL probes at five positions: C218 (**b**), C237 (**c**), C269 (**d**), C299 (**e**), C327 (**f**), the cyan line is the PRE effect calculated from 100 000 random $XRCC4_{CTR}$ conformations using Flexible Meccano (5). The residues are highlighted as in S12.

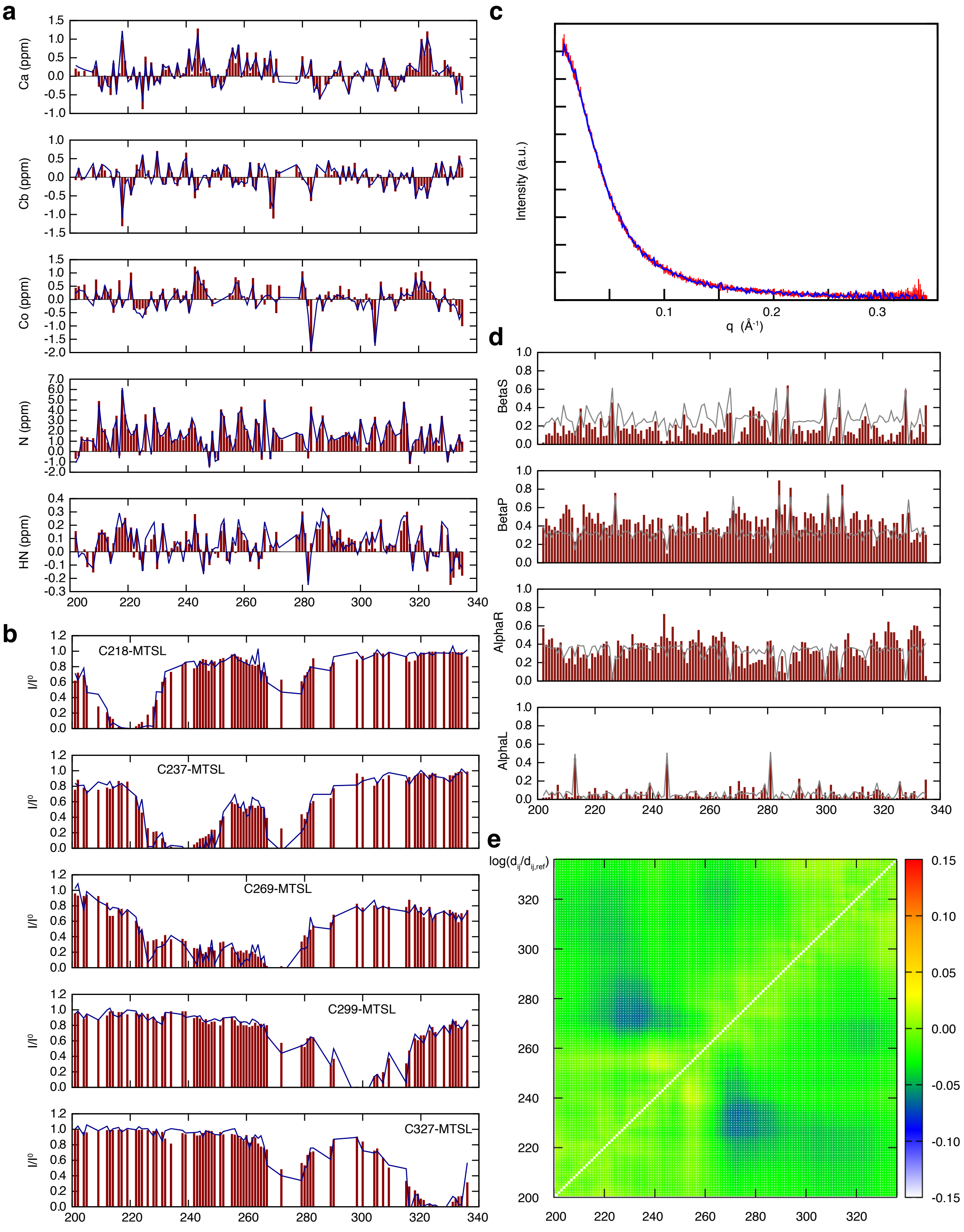


**Figure S16. Structural ensemble analysis of the XRCC4$_{CTR}$.** Comparison of the experimental (blue) and the ensemble back-calculated (red) data of (**a**) backbone chemical shifts, (**b**) PRE ratios and (**c**) SAXS curve. (**d**) Populations (red bars) of the secondary structure of the XRCC4$_{CTR}$ in the selected ensemble accordingly to the regions in the Ramachandran diagrams: β-sheet (BetaS), polyproline II (BetaP), α-helix (AlphaR), and left-handed helix (AlphaL). The gray lines indicate the statistical-coil distribution in the pool. (**e**) The log(10) matrix of the CA-CA distance ratio of the XRCC4$_{CTR}$ residues averaged over the selected ensemble and the statistical-coil pool. Red and blue colors indicate longer and shorter distances, respectively.

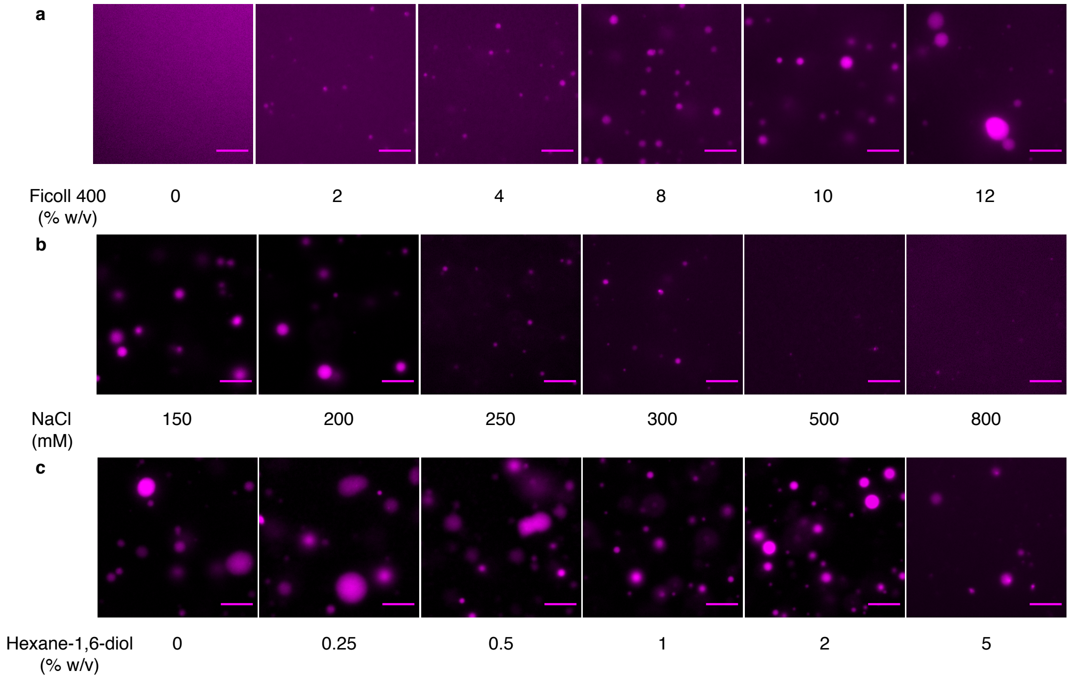


**Figure S17. The effects of additive components on LLPS of XLF and X4L4.** Fluorescence microscopy images of X4L4 (1% Cy3 labelling) and XLF with increasing concentrations of (**a**) Ficoll 400, (**b**) NaCl or (**c**) Hexane-1,6-diol. The concentration of X4L4 and XLF was fixed at 10 µM. The scale bar is 8 µm.

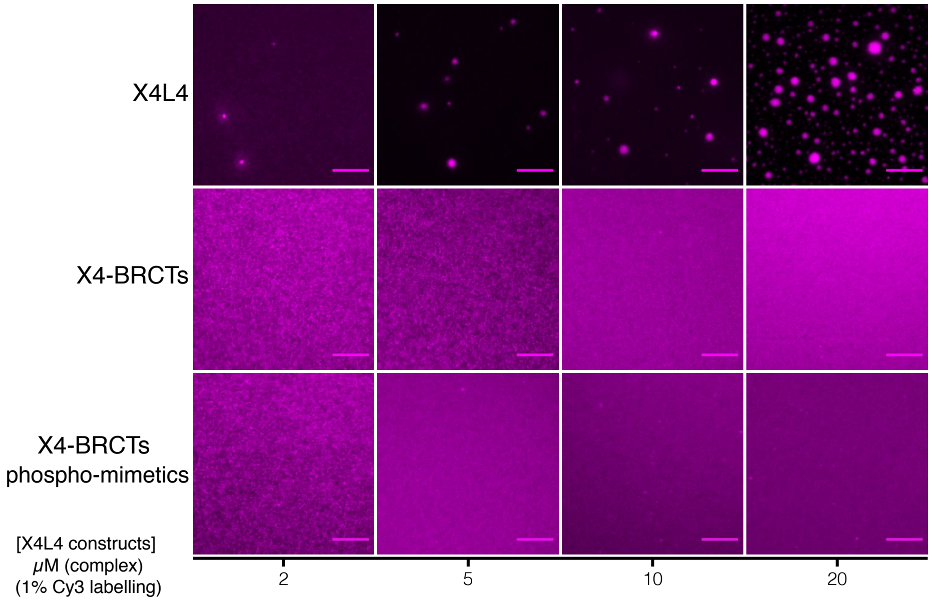


**Figure S18. X4L4 can undergo LLPS but not X4-BRTCs or X4-BRCTs phospho-mimetics.** Fluorescence microscopy images of 1% Cy3 labelling X4L4 constructs with increasing concentration. The scale bar is 8 μm.

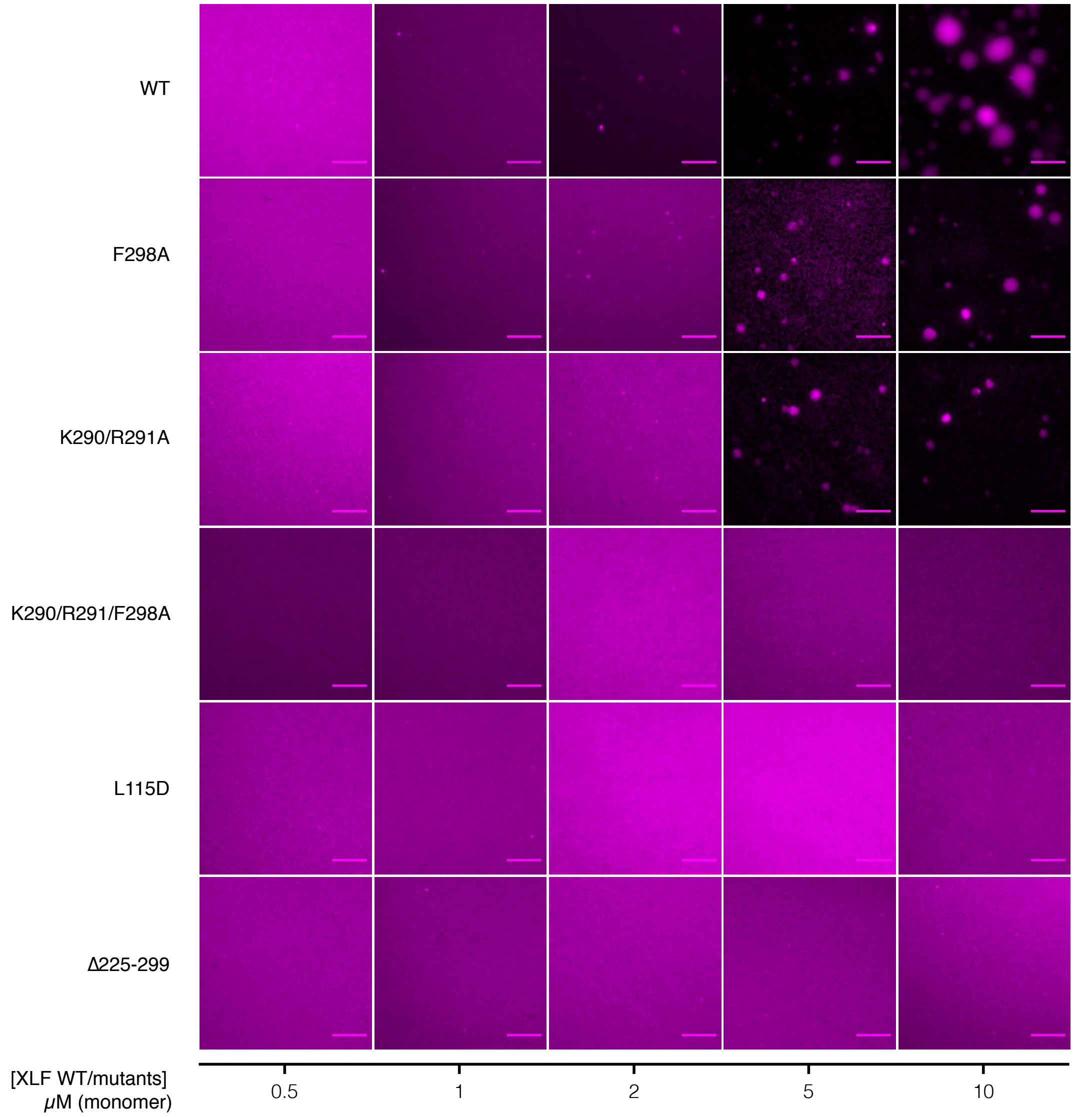


**Figure S19. The CTR of XLF is important for promoting LLPS with X4-BRTCs.** Fluorescence microscopy images of XRCC4-BRCTs with increasing concentrations of different XLF mutants. XLF variants were titrated into 10 μM 1% Cy3 labelling X4-BRCTs. The scale bar is 8 μm.

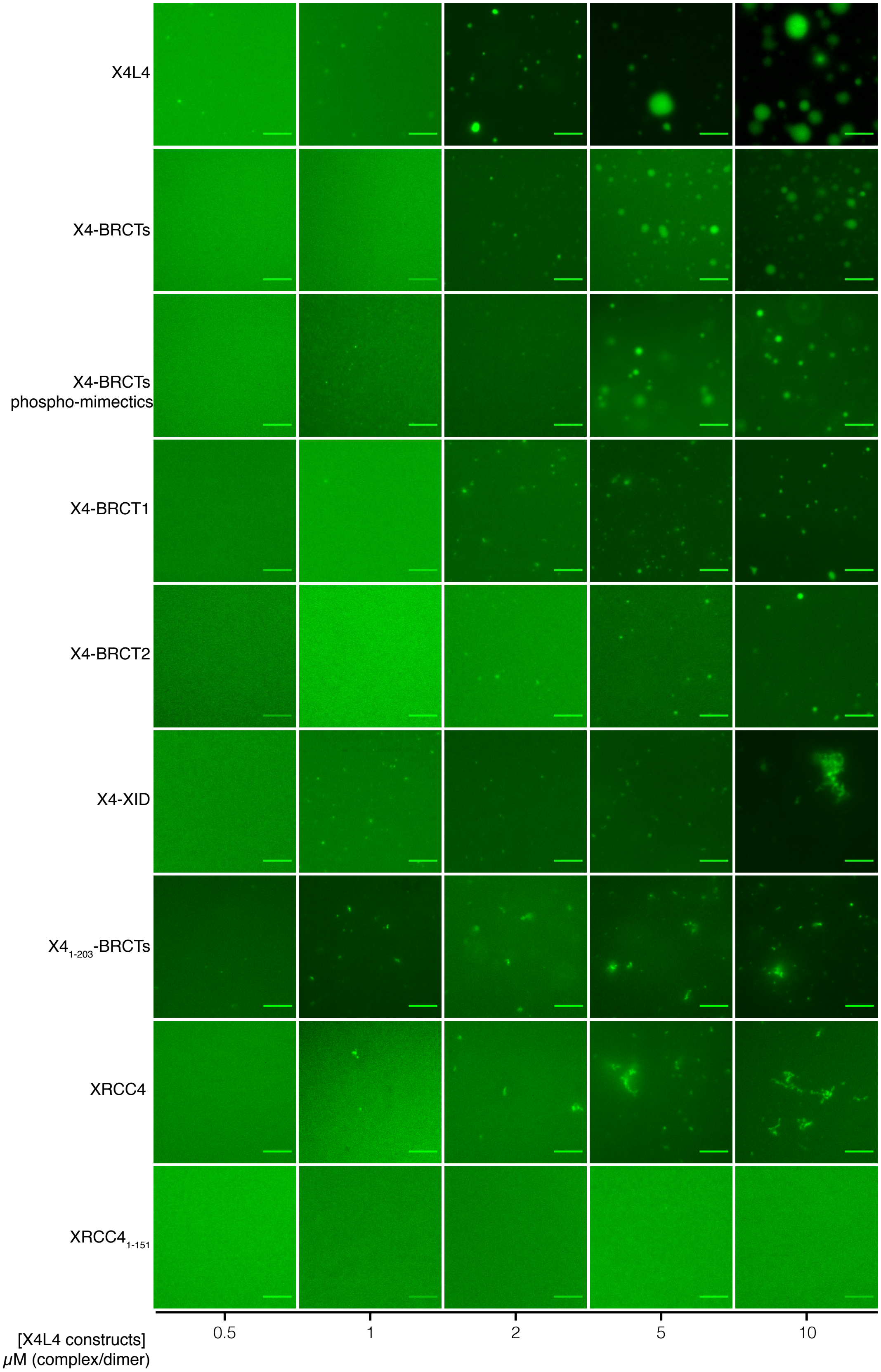


**Figure S20. The ability of different XRCC4 and LIG4 constructs undergo LLPS with XLF.** Fluorescence microscopy images of XLF (1% Fluorescein-labelled) with increasing concentrations of different XRCC4 and LIG4 constructs. XRCC4 and LIG4 variants were titrated into 10 μM (monomer) 1% Fluorescein-labelled XLF. The scale bar is 8 μm.

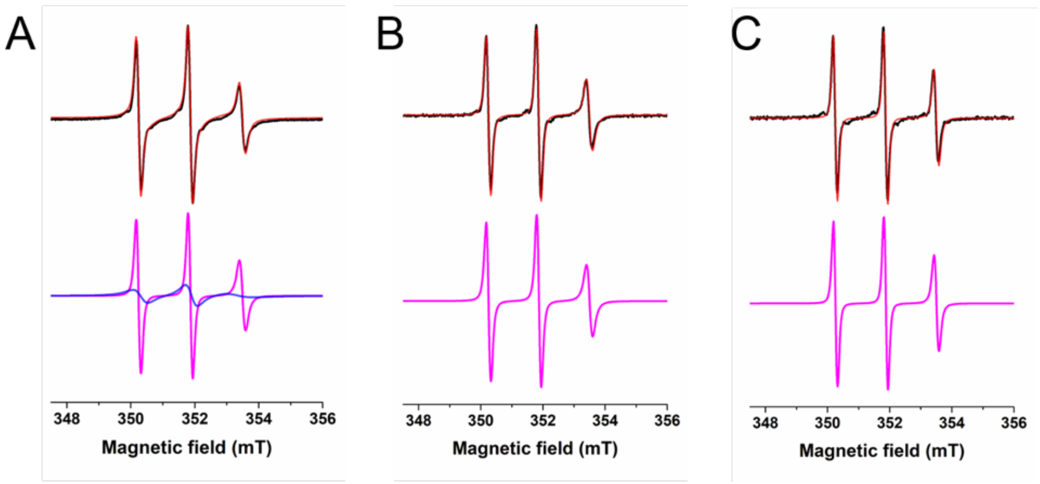


**Figure S21. EPR experiments of XRCC4.** Experimental (black) and simulated (red) CW-EPR spectra (X-band) for XRCC4 variants C218-MTSL (**a**), C299-MTSL (**b**) and C337-MTSL (**c**). Sharp (pink) and broad (blue) EPR component composing the whole EPR signals are reported under each spectrum. EPR simulations have been performed with SimLabel software (a GUI of EasySpin).

Continuous wave EPR spectroscopy (CW-EPR) was applied to probe the local dynamic of different XRCC4 regions within the CTR. The EPR spectra of different spin-labelled XRCC4 variants (**Fig. 3b**) show sharp spectral lineshapes, usually observed for IDRs and IDPs. Simulations of the EPR results (**Fig. S21**) revealed the presence of a component characterized by small values of correlational time ($\tau_C$) (**Table. S1, S2, S3**) for all the samples, suggesting a high flexibility of the CTR regions. However, the sharp component of the C218-MTSL spectrum represents 45% of the total EPR signal. Another component (55%, $\tau_C$= 2.00 ns) (**Fig. S21a**) defined by a broader lineshape indicates the presence of an additional dynamic state for this region. The EPR data confirm the intrinsically disordered nature of the $XRCC4_{CTR}$, as in agreement with the observation in our NMR investigation. In addition, no spectral variation was observed in the CW-EPR spectra of spin labelled XRCC4 variants in complex with BRCTs (**Fig. 3b**). These data reveal that CTR maintains its dynamic behaviour when BRCTs domain binds to the coiled-coil domain of XRCC4 and no major structural changes occur in the $XRCC4_{CTR}$ (**Fig. 3b**).

**Table S1.** Parameters used for the simulation of the experimental EPR spectrum of XRCC4 C218-MTSL using SimLabel software.

| **Component 1 (sharp component)** | | | | | |
|---|---|---|---|---|---|
| **gx** | 2.00873 | **Ax (mT)** | 0.50 | **component (%)** | 45 |
| **gy** | 2.00610 | **Ay (mT)** | 0.50 | $\tau_c$ **(nsec)** | 0.27 |
| **gz** | 2.00220 | **Az (mT)** | 3.90 | **lw gaussian (mT)** | 0.12 |
| **giso** | 2.00568 | **Aiso (mT)** | 1.63 | **lw lorenzian (mT)** | 0.04 |
| **Component 2 (broad component)** | | | | | |
| **gx** | 2.00779 | **Ax (mT)** | 0.50 | **component (%)** | 55 |
| **gy** | 2.00610 | **Ay (mT)** | 0.50 | $\tau_c$ **(nsec)** | 2.00 |
| **gz** | 2.00220 | **Az (mT)** | 3.90 | **lw gaussian (mT)** | 0.12 |
| **giso** | 2.00536 | **Aiso (mT)** | 1.63 | **lw lorenzian (mT)** | 0.04 |

**Table S2.** Parameters used for the simulation of the experimental EPR spectrum of XRCC4 C299-MTSL using SimLabel software.

| | | | | | |
|---|---|---|---|---|---|
| **gx** | 2.00853 | **Ax (mT)** | 0.50 | **component** (%) | 100 |
| **gy** | 2.00610 | **Ay (mT)** | 0.50 | $\tau_c$ **(nsec)** | 0.28 |
| **gz** | 2.00220 | **Az (mT)** | 3.90 | **lw gaussian (mT)** | 0.12 |
| **giso** | 2.00561 | **Aiso (mT**) | 1.63 | **lw lorenzian (mT)** | 0.04 |

**Table S3.** Parameters used for the simulation of the experimental EPR spectrum of XRCC4 C337-MTSL using SimLabel software.

| | | | | | |
|---|---|---|---|---|---|
| **gx** | 2.00853 | **Ax (mT)** | 0.50 | **component (%)** | 100 |
| **gy** | 2.00610 | **Ay (mT)** | 0.50 | $\tau_c$ **(nsec)** | 0.16 |
| **gz** | 2.00220 | **Az (mT)** | 3.90 | **lw gaussian (mT)** | 0.12 |
| **giso** | 2.00561 | **Aiso (mT)** | 1.63 | **lw lorenzian (mT)** | 0.04 |

**Table S4.** Plasmid constructs used in this work (refer to the supplementary sheet for the plasmid DNA and protein sequences).

| Lab collection number | Trivial name | Vector backbone (Selection) | Origin | Comments |
|---|---|---|---|---|
| pMM1610 | XLF vs1_pET-16b | pET-16b (AMP) | This work | produced in bacteria |
| pMM1611 | XLF New Cys74_pET-16b | pET-16b (AMP) | This work | produced in bacteria |
| pMM1612 | XLF New Cys188_pET-16b | pET-16b (AMP) | This work | produced in bacteria |
| pMM1613 | XLF 1-224_pET-16b | pET-16b (AMP) | This work | produced in bacteria |
| pMM1614 | XLF 1-261_pET-16b | pET-16b (AMP) | This work | produced in bacteria |
| pMM1615 | XLF vs1 L115D_pET-16b | pET-16b (AMP) | This work | produced in bacteria |
| pMM1616 | XLF vs1 L297E_pET-16b | pET-16b (AMP) | This work | produced in bacteria |
| pMM1617 | XLF vs1 F293A_pET-16b | pET-16b (AMP) | This work | produced in bacteria |
| pMM1618 | XLF vs1 T181E_pET-16b | pET-16b (AMP) | This work | produced in bacteria |
| pMM1619 | XLF vs1 K293A_pET-16b | pET-16b (AMP) | This work | produced in bacteria |
| pMM1620 | XLF vs1 K290R291AA_pET-16b | pET-16b (AMP) | This work | produced in bacteria |
| pMM1621 | XLF vs1 K290R291F298AAA_pET-16b | pET-16b (AMP) | This work | produced in bacteria |
| pMM1622 | XRCC4_pET-28a(+) | pET-28a(+) (KAN) | This work | produced in bacteria |
| pMM1623 | XRCC4 Cys0_pET-28a(+) | pET-28a(+) (KAN) | This work | produced in bacteria |
| pMM1624 | XRCC4 Cys218_pET-28a(+) | pET-28a(+) (KAN) | This work | produced in bacteria |
| pMM1625 | XRCC4 Cys299_pET-28a(+) | pET-28a(+) (KAN) | This work | produced in bacteria |
| pMM1626 | XRCC4 Cys337_pET-28a(+) | pET-28a(+) (KAN) | This work | produced in bacteria |
| pMM1627 | XRCC4 1-157_pET-28a(+) | pET-28a(+) (KAN) | This work | produced in bacteria |
| pMM1628 | XRCC4 1-203_pET-28a(+) | pET-28a(+) (KAN) | This work | produced in bacteria |
| pMM0365 | XRCC4 phospho-mimetic 8D | pET-28a(+) (KAN) | This work | DOI: 10.7554/eLife.22900 |
| pMM0903 | MBP G XID_pMBP-parallel1 | pMBP-parallel1 (AMP) | This work | produced in bacteria |
| pMM0905 | MBP I BRCT1-XID_pMBP-parallel1 | pMBP-parallel1 (AMP) | This work | produced in bacteria |
| pMM0907 | MBP K BRCT1_pMBP-parallel1 | pMBP-parallel1 (AMP) | This work | produced in bacteria |
| pMM0899 | MBP C BRCT2_pMBP-parallel1 | pMBP-parallel1 (AMP) | This work | produced in bacteria |
| pMM0901 | MBP E XID BRCT2_pMBP-parallel1 | pMBP-parallel1 (AMP) | This work | produced in bacteria |
| pMM0897 | MBP A BRCT1-XID-BRCT2_pMBP-parallel1 | pMBP-parallel1 (AMP) | This work | produced in bacteria |
| pMM1629 | XLF CTR 229-299 | pET-22b(+) (AMP) | This work | produced in bacteria |
| pMM1630 | XRCC4 CTR 200-336 | pMAL-c5x (AMP) | This work | produced in bacteria |
| pMM1631 | Ku80vWA 1-245 | pMAL-c5x (AMP) | This work | produced in bacteria |
| pMM1632 | MBP BRCTs Cys free | pMAL-c5x (AMP) | This work | produced in bacteria |
| pMM1633 | BRCT1 654-759 | pMAL-c5x (AMP) | This work | produced in bacteria |
| pMM1634 | ArtCTR | pGEX-6P-1 (AMP) | This work | produced in bacteria |

| | | | | |
|---|---|---|---|---|
| pMM1635 | Ku-SNAP = 10his-tev-Ku80/SNAP-Ku70 | pFL (AMP) | Gift from Jean-Baptiste Charbonnier, infected cell pellet provided | produced in insect cells |
| pMM0001 | X4LIG4 complex | pET28b(+) (KAN) | Gift from Murray Junop | produced in bacteria |
| pMM1603 | JS74 = pCAGGS-BSKX | pCAGGS (AMP) | Jeremy Stark Lab | PMID: 15485900 |
| pMM1604 | pX330 Cas9/sgRNA 7a for EJ7 | pX300 (AMP) | Jeremy Stark Lab | Addgene 113620 |
| pMM1605 | pX330 Cas9/sgRNA 7b for EJ7 | pX300 (AMP) | Jeremy Stark Lab | Addgene 113624 |
| pMM1577 | XRCC4 1-336 (isoform 1) | pCAGGS-BSKX (AMP) | This work | |
| pMM1579 | XLF 1-299 | pCAGGS-BSKX (AMP) | This work | |
| pMM1580 | XLF 1-251 | pCAGGS-BSKX (AMP) | This work | |
| pMM1589 | XRCC4 1-230 | pCAGGS-BSKX (AMP) | This work | |
| pMM1592 | XLF 1-299 K290R291F298AAA | pCAGGS-BSKX (AMP) | This work | |
| pMM1595 | XLF 1-299 L115A | pCAGGS-BSKX (AMP) | This work | |
| pMM1598 | XLF 1-299 L115A K290R291F298AAA | pCAGGS-BSKX (AMP) | This work | |
| pMM1599 | XLF 1-251 L115A | pCAGGS-BSKX (AMP) | This work | |
